# Natural loss of function of ephrin-B3 shapes spinal flight circuitry in birds

**DOI:** 10.1101/2021.01.29.428748

**Authors:** Baruch Haimson, Oren Meir, Reut Sudakevitz-Merzbach, Gerard Elberg, Samantha Friedrich, Peter V. Lovell, Sónia Paixão, Rüdiger Klein, Claudio V. Mello, Avihu Klar

## Abstract

Flight in birds evolved through patterning of the wings from forelimbs and transition from alternating gait to synchronous flapping. In mammals, the spinal midline guidance molecule ephrin-B3 instructs the wiring that enables limb alternation, and its deletion leads to synchronous hopping gait. Here we show that the ephrin-B3 protein in birds lacks several motifs present in other vertebrates, diminishing its affinity for the EphA4 receptor. The avian *ephrin-B3* gene lacks an enhancer that drives midline expression, and is missing in Galliformes. The morphology and wiring at brachial levels of the chick spinal cord resemble those of *ephrin-B3* null mice. Importantly, dorsal midline decussation, evident in the mutant mouse, is apparent at the chick brachial level, and is prevented by expression of exogenous *ephrin-B3* at the roof plate. Our findings support a role for loss of ephrin-B3 function in shaping the avian brachial spinal cord circuitry and facilitating synchronous wing flapping.

**Teaser:** Walking vs flying: Deciphering the organization and evolution of the neuronal network that controls wing flapping in birds.

## Introduction

A critical aspect of locomotion is left-right coordination. Limb alternation is the predominant gait mode in some amphibians (salamanders), reptiles and terrestrial mammals, while synchronous movements are manifested in the legs of other amphibians (frogs) and some marsupials, and in the wings of bats and birds. Based on classical electrophysiological studies in cats and rodents, and genetic manipulations of cell fate and axonal trajectories in mice, a modified “half-center oscillator” model has been proposed that explains limb alternations in mammals (*1, 2*). The core elements are mutually inhibitory neuronal centers on each side of the spinal cord that produce oscillatory excitatory output. This output is simultaneously transmitted to interneurons that innervate motoneurons, termed pre-motoneurons (pre-MNs). These include ipsilateral excitatory pre-MNs that activate motor activity on one side, and commissural inhibitory pre-MNs that inhibit motor output on the other side (Fig. 1A) (*3–6*).

**Figure 1:**
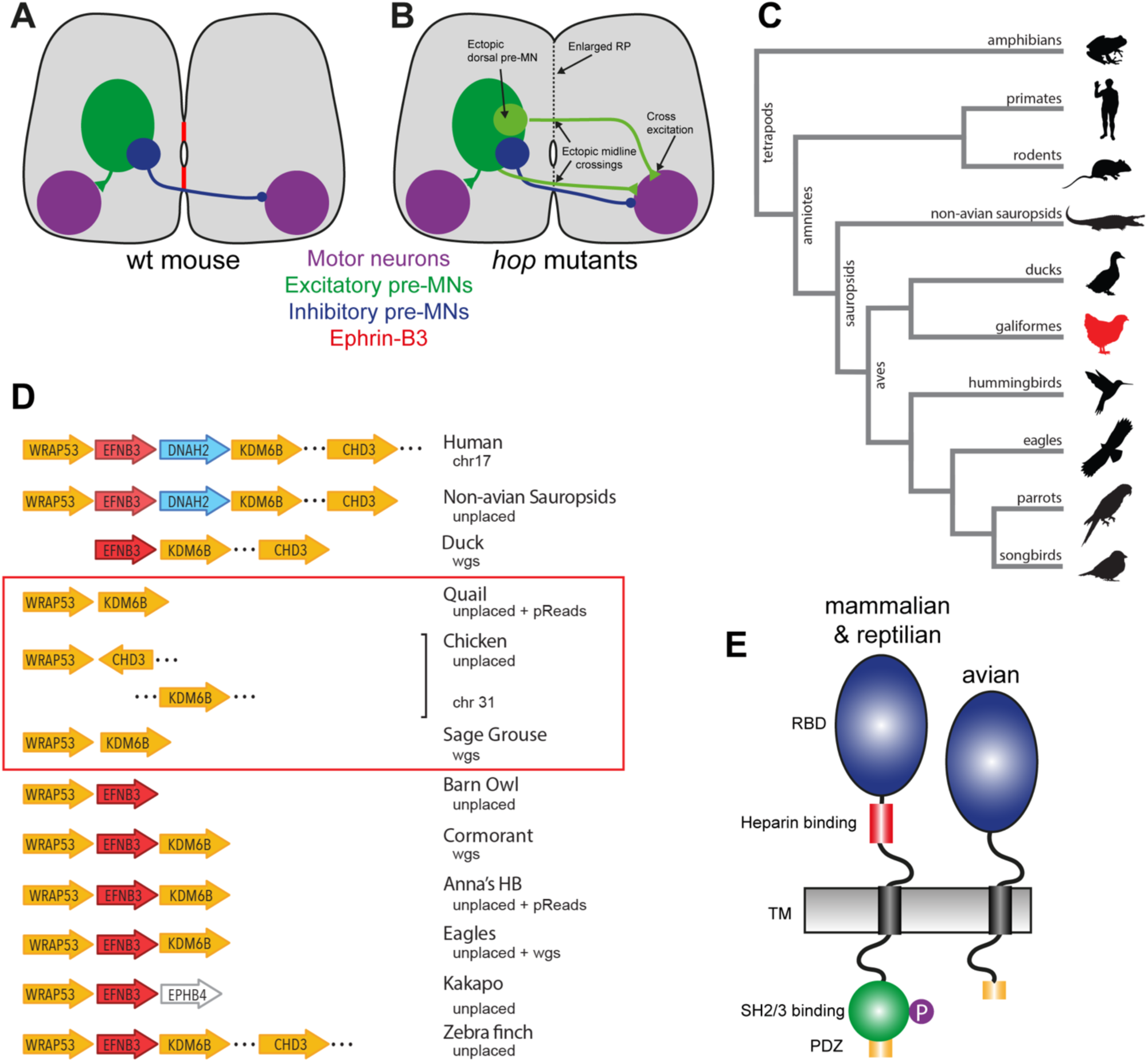
Spinal gait circuitry and EFNB3 genomics. **A,B**. A scheme illustrating the morphology and wiring of the spinal cord in wt mouse (A) and in ephrin-B3/EphA4/12-chimaerin mice collectively referred to as *hop* mutants (B); pre-MN: pre- motoneurons, RP: roof plate. **C.** Simplified cladogram depicting the relations across representative organisms examined in this study. Branch lengths are arbitrary and not calibrated for time, red means EFNB3 is absent in chicken. **D.** Comparison of EFNB3 (red) synteny across organisms where synteny could be examined, with chromosomal location or data source indicated. EFNB3 is absent in Galliformes (red square) and DNAH2 (blue) is absent in birds. Non-avian sauropsids: lizard and alligator; wgs: whole-genome shotgun contigs; HB: hummingbird. **E.** Schematic structure of the predicted mouse and avian ephrin-B3 proteins. RBD: receptor binding domain; TM: transmembrane domain.

The axon guidance molecule ephrin-B3 and its cognate receptor EphA4 are key regulators of alternating gait. In this family, ligand binding to the receptor typically results in axonal repulsion (forward signaling), but ephrin receptors can also act as ligands for ephrins (reverse signaling), prompting repulsion or adhesion (*7*). In wt mice, ephrin-B3 serves as a midline repulsive cue, preventing EphA4-expressing axons of excitatory pre-MNs from aberrantly crossing the ventral and dorsal midlines (Fig. 1A) (*8–12*). Transition from left-right alternation to synchrony occurs in mice mutated in the genes encoding ephrin-B3, EphA4, or the downstream signaling molecule α2- chimaerin, referred to as *hop* mutants. These mice show ectopic commissural projections of otherwise ipsilaterally-projecting excitatory pre-MNs (Fig. 1B) and walk with synchronous movements of the hind- and forelimbs, producing a hopping gait. Furthermore, impairment of the neuroepithelial organization of the roof plate (RP), which leads to partial absence of ephrin-B3 expression, results in rabbit-like hopping (*13, 14*).

Birds have evolved from dinosaurs that use alternating gait (*15, 16*). Their adaptation for flight includes substantial structural specializations of the forelimb and of the spinal cord circuits that enable synchronous limb movements. Classical transplantation experiments in chick support a role for neuronal networks at the lumbar and brachial spinal levels in the control of the locomotion pattern of the hind- and forelimbs, respectively. In chicks where brachial segments are surgically replaced by lumbosacral segments, the movement of the wings become alternation, whereas coordinated flapping movements are not observed (*17*). In the complementary experiment, transplantation of brachial segments to the lumbar cord result in synchronous, hopping-like leg movements (*18*). These findings support the hypothesis that local circuits in the lumbar and brachial spinal cord instruct level-specific alternate or synchronous movements, respectively.

Flight requires spinal circuits that enable the bilateral coordination of forelimb movements. It thus seems reasonable to hypothesize that the emergence of synchronous wing flapping in birds involved a transition from ipsilateral excitatory/contralateral inhibitory inputs to a pattern of bilateral co-excitatory input to wing motoneurons. Such a transition likely required changes in the pre-motor spinal wiring. Notably, mutations in the midline guidance molecules Netrin1 or ephrin- B3 in mice lead, respectively, to a reduction in cross inhibition and an increase in cross excitation (*9, 19, 20*). Netrins were initially described in chick, and have well characterized activity and roles in commissural axonal guidance (*21, 22*). However, little is known about ephrin-B3 in chick, and more generally in birds.

To explore a possible role of ephrin-B3 in locomotion patterns in birds, we have conducted a set of genomic, molecular and neuroanatomical studies, using both chick and zebra finch as experimental organisms. We found an ephrin-B3 gene ortholog (EFNB3) in several bird species and lineages, however their open reading frames (ORFs) were considerably shorter, and some domains critical for forward and reverse signaling were absent or heavily modified compared to non-avian tetrapods. We also observed marked loss of dorsal midline ephrin-B3 expression in the brachial spinal cord of zebra finch, likely due to an avian loss of a critical enhancer element. An EFNB3 ortholog could not be found in chicken and related species, suggesting a gene loss in Galliformes. Spinal cord interneurons in chick showed extensive dorsal midline crossing at brachial levels, similar to ephrin-B3 null mice. In contrast, we observed little dorsal crossing at lumbar levels due to the presence of the glycogen body, which serves as a physical barrier to neurite crossing. The crossing at brachial levels was largely prevented by dorsal midline expression of mouse ephrin-B3. These findings are consistent with a role of ephrin-related mechanisms in spinal wiring patterns that may subserve limb alternation. They also support the notion that diminished barriers to dorsal midline crossing may have favored synchronous limb movements and facilitated the origin and/or maintenance of coordinated flight in birds.

## Results

### EFNB3 gene in birds

We have conducted an extensive and thorough analysis of the genomic region containing EFNB3 and syntenic genes in birds and non-avian outgroups. Our approach required extensive curation of gene predictions in annotated databases, permissive BLAST searches of genomes, WGS databases and p-reads (from Pacbio assemblies) using avian EFNB3 and syntenic gene queries, and RefSeq alignment searches of unplaced genomic scaffolds and reads (details in Methods). We found EFNB3 in a broad range of bird species and orders, including several songbirds, some eagles, an owl, a few cormorants, as well as the seriema, a hummingbird, a ruff, a crane, a stork, and some Anseriformes (duck and goose). WRAP53 was immediately upstream of EFNB3 in all birds where synteny could be examined and in all non-avian organisms studied, including mammals, non-avian sauropsids and amphibians (examples in Fig. 1C,D), providing strong support for EFNB3 orthology. A gene prediction was present in only few species, whereas in several others EFNB3 was partial, possibly due to incomplete sequences in Illumina assemblies.

Notably, we could not identify EFNB3 in any Galliformes (Fig. 1D, red square), even in species with high quality Pacbio assemblies and no local gaps. In chicken, the genes immediately syntenic to EFNB3 in other birds (WRAP53 and KDM6B) were on different chromosomes, suggestive of a local inter-chromosomal rearrangement in this species, with no trace of EFNB3 (Fig. 1D, chicken). EFNB3 was also absent in the chicken WGS and Pacbio p-reads, and in numerous RNAseq databases. In quail, a WRAP53 prediction and an unannotated KDM6B (found by BLAST) were found on the same incomplete scaffold. Several Pacbio p-reads from the current assembly (*23*) bridged the local gaps but revealed no trace of EFNB3 (Fig. 1D, quail). EFNB3 was also not detected in the assemblies or WGS of other Galliformes (turkey, pheasant, guinea fowl), even though the syntenic genes were sometimes present in unplaced scaffolds (not shown). A scaffold in the unannotated sage grouse genome contained traces of WRAP53 and KDM6B but no evidence of EFNB3 sequences (Fig. 1D, sage grouse). We also could not find EFNB3 in ratites, but the data were too sparse to conclusively prove or disprove an EFNB3 loss in this basal group

Importantly, in most birds where EFNB3 synteny could be verified KDM6B was immediately downstream, whereas in non-avian sauropsids and mammals DNAH2 was interposed between EFNB3 and KDM6B (Fig. 1D, blue). Given there were no gaps in several avian assemblies between EFNB3 and KDM6B, this observation supports an avian-specific loss of DNAH2, as previously indicated (Lovell et al., 2014). As we show later, this finding has important implications for EFNB3 expression regulation. The exception among birds was the kakapo, where the receptor gene EphB4 was downstream of EFNB3, the only such case we are aware of. In sum, the EFNB3 gene is clearly present in a broad range of Neoaves, as well as in Anseriformes, but seems absent in Galliformes as a group (Fig. 1C,D). We suggest this is possibly due to a loss that occurred after their split from Anseriformes, an estimated 46 million years ago (*16*), but prior to their later diversification.

### EFNB3 gene and protein structure in songbirds

Songbirds were the avian group with the most complete information on EFNB3. In zebra finches, EFNB3 is largely conserved compared to non-avian tetrapods, consisting of 5 exons and a large 3’- UTR (Fig. S1A), but predicted avian ephrin-B3 proteins are shorter compared to amphibian, reptilian and mammalian EFNB3 (247-267 vs 332-340 aminoacid residues; Fig. S2). The intracellular domain is shorter in birds, and while the PDZ motif is intact, the entire SH2/3 binding motif, which is required for reverse signaling (*24, 25*), as well as the tyrosine residues that are phosphorylated upon activation of the reverse signaling in other organisms, are missing (Fig. 1E; Fig. S2), suggesting that reverse signaling is impaired. Birds also lack an extracellular heparin binding motif within the region juxtaposed to the transmembrane (TM) domain (*26, 27*), and several residues in the tetramerization interface within the receptor binding domain (RBD), required for normal signaling through the formation of receptor-ligand (Eph–ephrin) complexes (*28*) (Fig. 1E; Fig. S2). The region in the dimerization interface where an L to P (L111P) substitution leads to a hopping mouse phenotype (*29*) is conserved in birds and shows other substitutions (DLDL_108-111_QRDV) compared to mammals (Fig. S2). Unlike ephrin-B3, the predicted avian ephrin-B1 and ephrin-B2 proteins are highly similar to the mammalian proteins in terms of size, sequence and sub-domain partitioning.

To test whether these structural differences affect cell trafficking of the avian ephrin-B3, we expressed an amino-myc tagged isoform of the mouse and zebra finch genes in COS-7 cells. Surface staining revealed that both proteins are presented on the cell membrane (Fig. 2A). Next, we used an assay where a chimeric protein composed of the extracellular domain of chicken EphA4 fused to placental alkaline phosphatase is tested for binding to different EFNB3 orthologs expressed in transiently transfected COS-7 cells, noting that chick and the zebra finch EphA4 proteins share 98.4% identity. We found that chicken EphA4 binds mouse EFNB3, but not zebra finch EFNB3 (Fig. 2B,C).

**Figure. 2:**
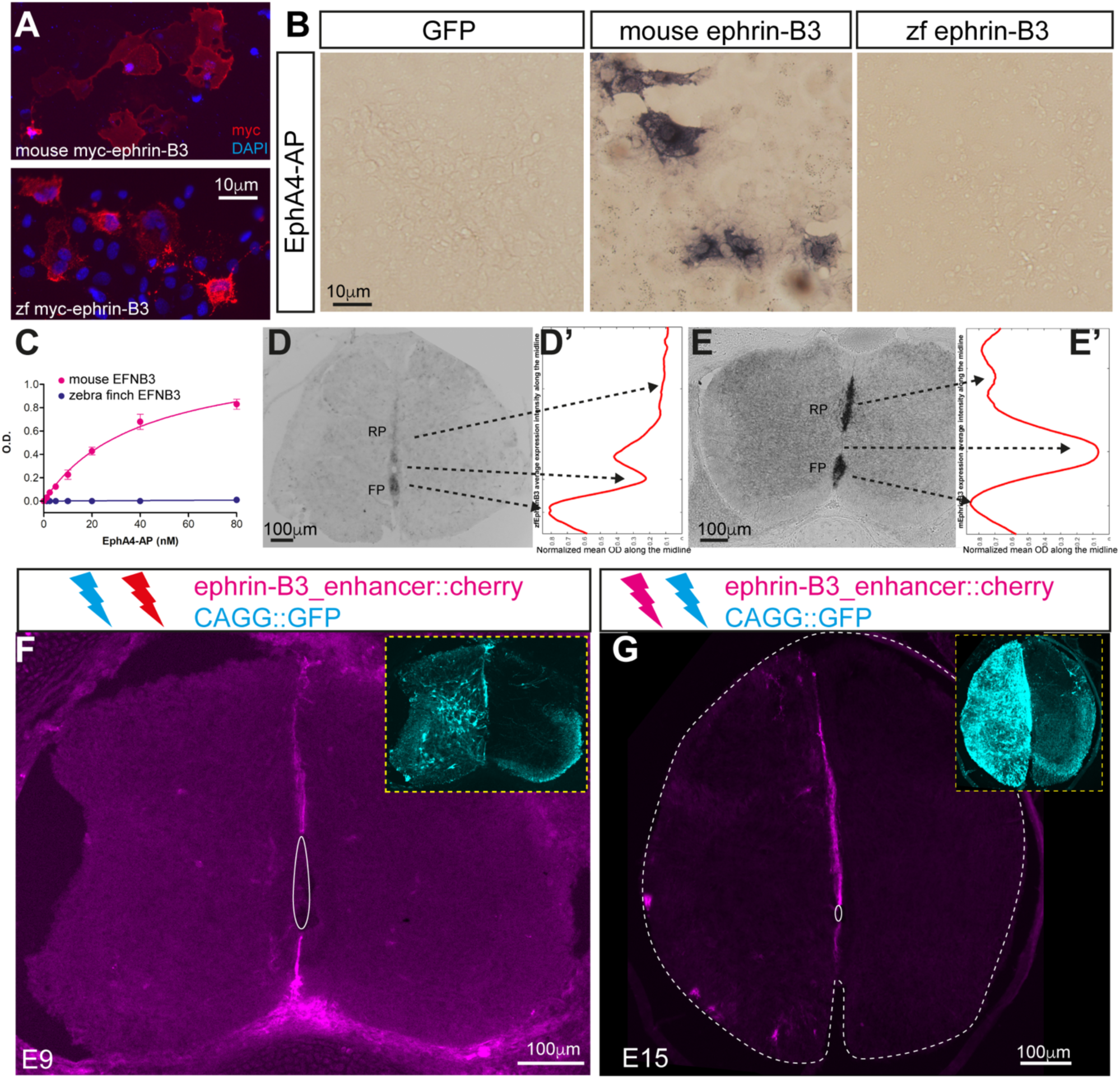
Zebra finch ephrin-B3 protein and mRNA expression. **A.** The mouse and zebra finch proteins are presented on the cell membrane. Anti myc antibody (9E10) was used for surface staining of COS-7 cells transiently transfected with amino-terminal myc-tagged ephrin-B3 isoforms. The sequence of the full length zebra finch ephrin-B3 transcript was reconstructed from RNA-seq reads using the Trinity platform (*59*) (Supp. Fig. S1). **B.** Binding assay shows EphA4-AP binding to mouse ephrin-B3 protein but not to GFP or to zebra finch ephrin-B3. **C.** Representative EphA4-AP/ephrin-B3 binding curves assessed by AP activity in a colometric assay. Apparent KD value for mouse ephrin-B3 was 40.15 nM (n=3).**D.** *In situ* hybridization for ephrin-B3 mRNA in P0 zebra finch spinal cord. High expression is evident in the floor plate (FP), but declines along the roof plate (RP). **D’.** Normalized optical density (OD) profile of zebra finch ephrin-B3 midline expression; shown is the means from 58 cross sections. **E.** *In situ* hybridization for ephrin-B3 mRNA in E15.5 mouse spinal cord. High expression is evident in both the FP and RP. **E’.** Normalized optical density (OD) profile of mouse ephrin-B3 midline expression; shown is the means from 160 cross sections. **F,G.** Analysis of the activity of the ephrin-B3 enhancer element, at E9 (F) and E15 (G). The cherry reporter gene was cloned downstream to the putative enhancer and electroporated into the chick spinal cord. GFP, under the control of the ubiquitous CAGG enhancer-promoter, was co-electroporated. The reporter is detected in both FP and RP (F,G), similar to the pattern of ephrin-B3 expression in mice, while GFP is expressed ubiquitously.

### Expression of ephrin-B3 in zebra finch

The presence of EFNB3 gene in songbirds prompted us to examine its expression pattern in the zebra finch spinal cord. *In situ* hybridization with antisense riboprobes revealed ephrin-B3 mRNA expression in both the floor plate (FP) and roof plate (RP) of the brachial spinal cord in P0 zebra finches, with apparent lower expression in the RP (Fig. 2D). Quantification confirmed that ephrin- B3 expression is higher in the FP and declines along the RP in the ventral-to-dorsal direction (Fig. 2D’). This contrasted with mouse at a comparable developmental stage, where expression is similarly high in both the FP and RP (Fig. 2E), with no dorsal decline (Fig. 2E’).

We reasoned that the modified ephrin-B3 expression seen in zebra finch compared to mouse might relate to changes in regulatory elements. Using UCSC’s genome browser, we searched for enhancer hallmarks in mouse EFNB3 and its flanking genes. We found that mouse DNAH2, located immediately downstream of EFNB3 (Fig. 1D, blue), contains several features consistent with presence of an enhancer element in its 1^st^ intron (Fig. S3). Because the DNAH2 gene is not present in birds (Fig. 1D blue; Lovell et al., 2014), birds may lack an important EFNB3 enhancer element. To test whether this DNAH2-embedded region would be sufficient to confer midline RP expression in the spinal cord of a bird species, a reporter gene cloned downstream of the putative enhancer element was electroporated into the chick spinal cord at E3. Expression of the reporter in both the FP and RP was evident at E9 and E15 (Fig. 2F,G), similar to mouse where endogenous ephrin-B3 midline expression occurs during embryogenesis and early post gestation (E15 chick is equivalent to mouse P2), and contrasting with the more ubiquitous expression of the co-electroporated GFP under a ubiquitous promoter (Fig. 2F,G, insets). Notably, *DNAH2* is missing in all bird species examined to date but it is present in reptiles (Fig. 1D; (*30*)), suggesting the loss of the EFNB3 regulatory element is specific to the avian lineage.

### Expression of ephrin-B family ligands and EphA4 in chick spinal cord

We next used *in situ* hybridization to examine expression of ephrin-B1 and -B2 at the chick brachial cord, as both can mediate axonal repulsion through the EphA4 receptor and could thus potentially compensate for an EFNB3 loss. Both ephrin-B1 and B2 were expressed in the FP and RP at E10 (Fig. 3A,C). At E16, when the spinal circuitry elicits simultaneous wing flapping (*31*), ephrin- B1 expression persisted in the FP but dorsally it became restricted to the ventral aspect of the RP (Fig. 3B), whereas ephrin-B2 expression was detected only at the ventral tip of the RP (Fig. 3D). Expression of ephrin-B genes in chick spinal cord is thus largely restricted to the FP, low in ventral RP, and absent in the dorsal RP.

**Figure 3:**
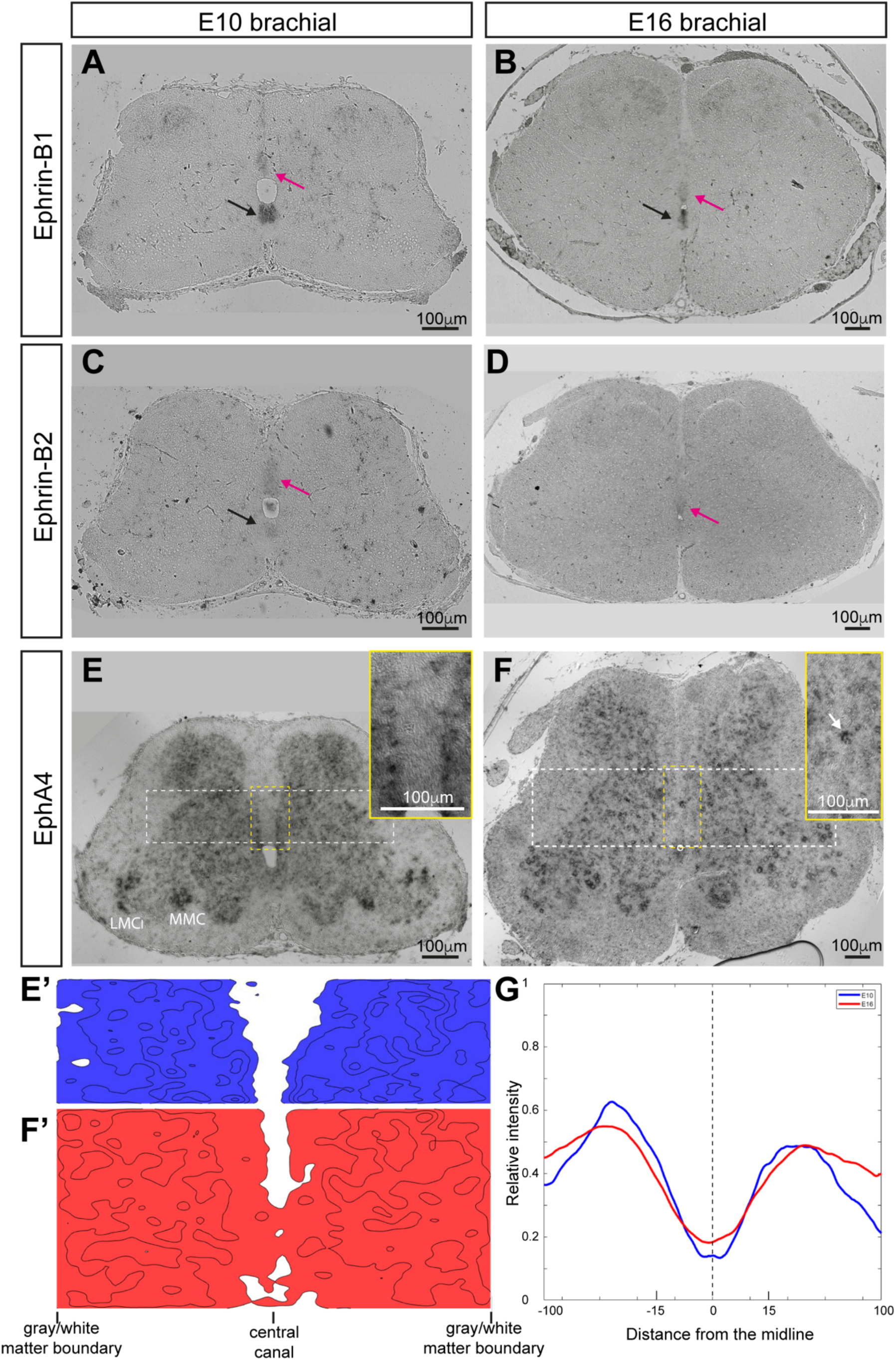
Expression of ephrin-B1, ephrin-B2 and EphA4 mRNAs in the chick spinal cord. *In situ* hybridization for ephrin-B1 (A,B), ephrin-B2 (C,D) and EphA4 (E,F) mRNAs at the brachial level in E10 (A,C,E) and E16 (B,D,F) cords. The floor plate (FP) is indicated by black arrows and the roof plate (RP) by magenta arrows. **A,B**. Ephrin-B1 at E10 is high in the FP and low in the RP at E16 it is high in the FP and low in ventral RP. **C,D.** Ephrin-B2 at E10 is low in the FP and RP; at E16 it is low in ventral RP. **E,F.** EphA4 expression at E10 is widespread in the gray matter, with higher levels in scattered interneurons the lateral subdivision of the lateral motor columns (LMCl) and the medial motor columns (MMC). At E16, expression is detected in scattered interneurons and motoneurons. Interneurons invading the dorsal midline are apparent (white arrow in inset in F). Insets in E and F are enlargements of the areas boxed in dashed yellow. **E’,F’.** Cell density distribution, extracted from 10 cross sections from the regions flanking the dorsal midline at E10 and E16, respectively (indicated by white dashed boxes in E and F). **G.** Summation of cell *in situ* hybridization relative intensity in the area indicated in E’ (E10, blue) and F’ (E16, red). The area along the X was subdivided to 100 epochs. N=10 sections for each experiment. Values are significantly different around the dorsal midline (spanning from -15 to 15) p<0.0001, t-test, see Materials and methods and supplementary statistics).

Next, we studied the expression of the receptor EphA4. In mouse, interneurons that express EphA4 are excluded from the dorsal midline, but many of these cells invade the dorsal midline in *EphA4* and *ephrin-B3* null mutants (*32*). In the chick, expression is detected throughout the spinal cord gray matter at E10, with high levels in scattered interneurons, in the lateral subdivision of the lateral motor column (LMCl), and in the medial motor column (MMC), but absent in the dorsal midline (Fig. 3E). At E16, EphA4 shows high expression in discrete cells distributed all over the gray matter, including adjacent to and within the dorsal midline (Fig. 3F, white arrow in inset).

When we assigned x-y coordinates to EphA4-expressing cells, we observed their midline invasion at E16 compared to E10 (Fig. 3E’,F’,G). Thus, the presence of EphA4-expressing neurons in the dorsal midline correlates with the temporal reduction of ephrin-B1-2 levels. Taken together with the lack of ephrin-B3, we conclude that a molecular barrier that prevents midline crossing is missing in the chick dorsal spinal cord. This contrasts markedly with mouse, where midline expression of ephrin-B3 persists at high levels both ventrally and dorsally at post gestation stages (*9*).

### Extended roof plate of the chick brachial cord

To further investigate how the natural loss of EFNB3 might have affected the organization of the avian spinal cord, we examined several parameters that differ significantly between wt and *hop* mutants, namely the size and axonal decussation at the RP, and the somata distribution and neurotransmitter phenotype of commissural pre-MNs (Figs. 4-6).

**Figure 4:**
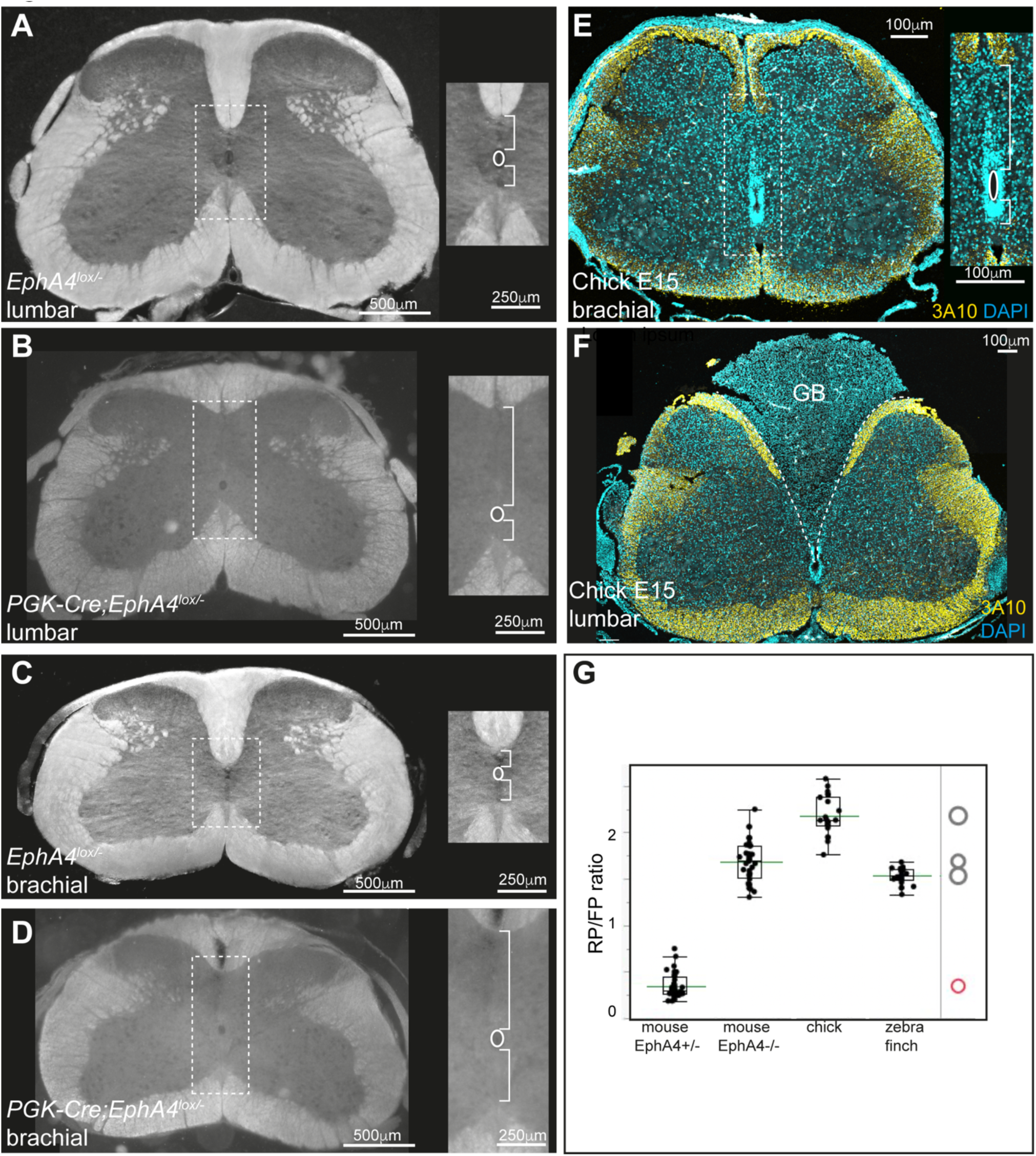
Spinal cord morphology in *hop* mutants and chick show enlarged brachial RPs. **A-D**, Dark field microphotographs of transverse sections of adult mouse spinal cord at lumbar and brachial levels. Compared are control heterozygous (EphA4^lx/lx-^; A,C) and homozygous null mice (EphA4^lx/lx-^/PGK-Cre^+^; B,D) (*32*). Right insets (enlargements of the boxed areas) depict the central canal and the RP and FP. **E,F.** Transverse sections at the brachial (E) and lumbar (F) levels of E15 chick spinal cord. The white matter is labelled with the anti-neurofilament antibody 3A10, the gray matter with nuclear DAPI staining. Right inset (in E) is an enlargement of the boxed area. **G.** Box plot representation of the roof plate (RP) to floor plate (FP) length ratios at the brachial level in adult EphA4+/- mouse (N= 3, 36 cross sections), adult EphA4-/- mouse (N= 3, 36 sections), E15 chick (N= 2, 22 sections) and P0 zebra finch (N= 1, 25 sections). The difference between the heterozygous mouse, chick and zebra finch groups and the control homozygous null mouse group is statistically significantly (See supplementary statistics). RP: roof plate; FP: floor plate.

In wt mice, the length of the FP was relatively constant along the longitudinal axis, but the RP was substantially shorter at limb levels, resulting in low RP:FP ratios at those levels (Fig. S4). Thus, the RP:FP ratio can serve as a reasonable proxy for the size of the RP. The length of the RP at limb levels was markedly enlarged in EphA4 knockout mice (Fig. 4B,D) in comparison to control mice (Fig. 4A,C), with a significantly higher RP:FP ratio in the homozygous mutant (1.68±0.22, mean of RP:FP ratios ± SD) than in control mice (0.34±0.13). In birds, an elongated RP was evident at brachial levels in E15 chick (Fig. 4E) and P8 zebra finch (not shown), with RP:FP ratios of 2.18±0.2 and 1.53±0.08, respectively, thus comparable to EphA4 knockout mice (Fig. 4G). Hence, the morphology of the avian brachial RP is similar to that of *hop* mutants. At the lumbar sciatic level, the dorsal midline in birds is occupied by an ovoid gelatinous mass - the glycogen body, an avian specific organ (GB; Fig. 4F) (*33–35*). These features differ from those in alternating limbed reptiles (lizard, turtle) where the RP is short, as in wt mice (*36–38*)

### Patterns of axonal midline crossing in the chick spinal cord

Axon fibers that cross the RP in rodents originate mostly from sensory neurons and inhibitory interneurons (*39, 40*). To further study the extent of dorsal midline crossing in the chick embryonic spinal cord, we examined immunoreactivity for neurofilament (3A10) and for sensory axons (TAG1) at E15 and E17. We observed neurofilament-positive axons crossing the brachial FP in a tight bundle (Fig. 5A, yellow arrows in insets), whereas those crossing the dorsal midline were less fasciculated and sparsely distributed along the dorsal RP (Fig. 5A, white arrows in insets). Quantification showed that 44.8±23% of midline crossing axons at the brachial level were in the RP (Fig. 5G, brachial). TAG1-expressing sensory axons did not appear to cross the midline at this level (Fig. 5B). To further interrogate midline axonal crossing, a membrane-tethered GFP was electroporated at E3 into one side of the spinal cord. At E15, GFP-labeled axons crossing the ventral and dorsal midline were evident (Fig. 5C, yellow and white arrows in insets). Interestingly, a few labeled cells were apparent on the contralateral side dorsally, suggesting that some cells may migrate through the RP at the brachial level (Fig. 5C, white arrowheads in insets).

**Figure 5:**
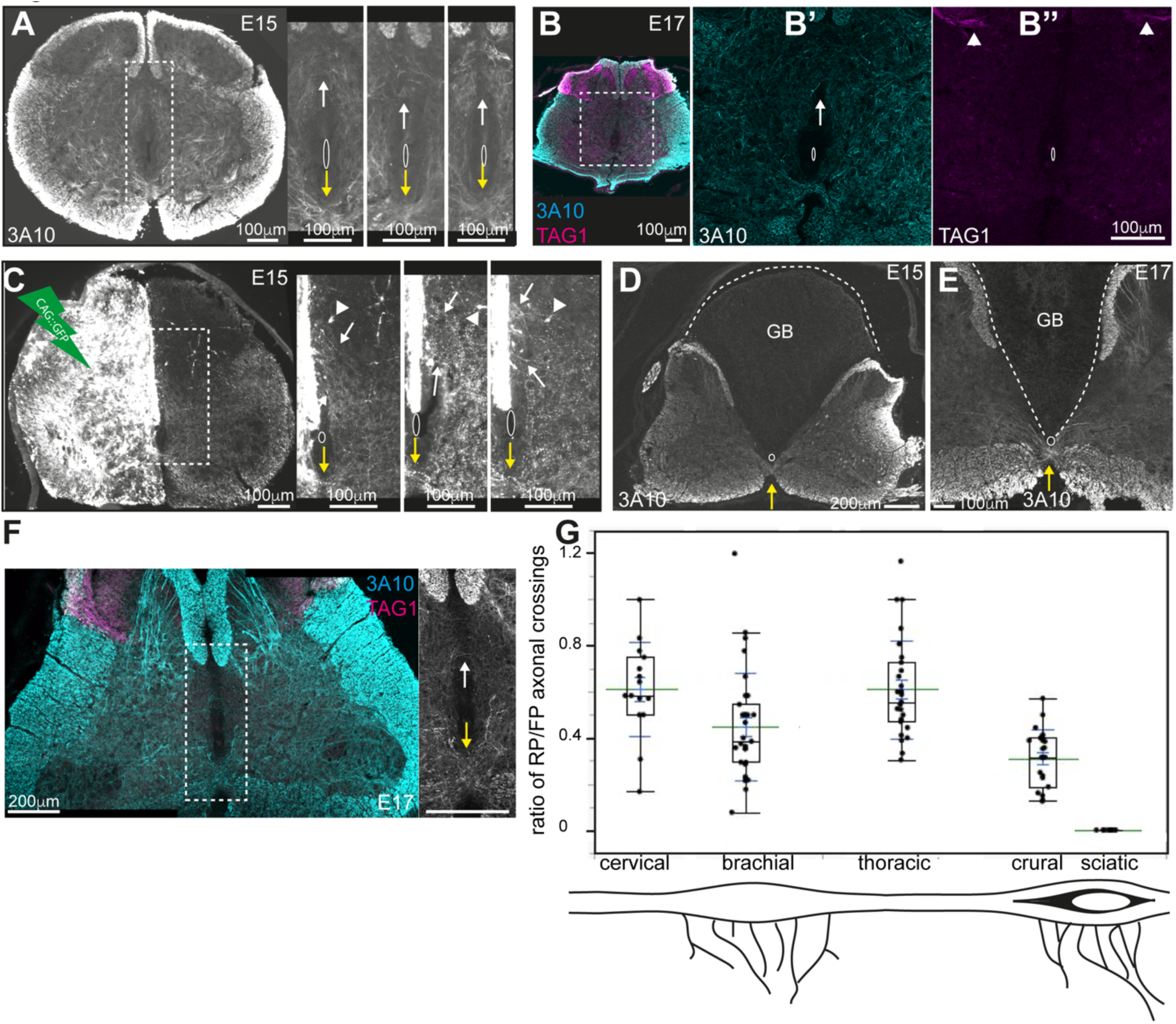
Midline crossing at the brachial and lumbar levels of chick spinal cord. **A.** Dorsal midline crossing at brachial level in E15 chick demonstrated by neurofilament (3A10) immunostaining in transverse sections. Detail views of three examples in insets, left inset is an enlargement of the boxed area in A. Axons cross at the FP (yellow arrows) and the dorsal part of the RP (white arrows). **B.** Dorsal midline crossing at brachial level in E17 chick; the boxed area is enlarged in B’ and B’’. Anti-3A10 labels all axons, and anti-TAG1 labels sensory axons (*61*). **C.** Dorsal midline crossing is evident following unilateral expression of a reporter gene. GFP under the control of the CAGG enhancer/promoter construct was electroporated at E3 into the left side of the spinal cord. At E15, midline-crossing axons are evident in the brachial FP (yellow arrows) and RP (white arrows) . Detail views of three examples in insets, left inset is an enlargement of the boxed area in C. **D,E.** Transverse sections at the sciatic plexus level at E15 (D) and E17 (E). Axons cross only at the FP (yellow arrows). The glycogen body (GB) lacks axonal labelling. **F.** Transverse section at the crural plexus level at E17. Right inset is an enlargement of the boxed area in F. Axons cross at the FP (yellow arrow) and RP (white arrow). **G.** Box plot of the ratio between axons crossing the RP and the FP at different axial levels in E15 chick. Numbers of midline-crossing axons were scored at cervical (15 sections), brachial (31 sections), thoraxic (27 sections), lumbar crural (23 sections) and lumbar sciatic (14 sections) levels of one embryo. At the sciatic level axons did not cross the glycogen body. (See supplementary statistics). RP: roof plate; FP: floor plate.

**Figure 6:**
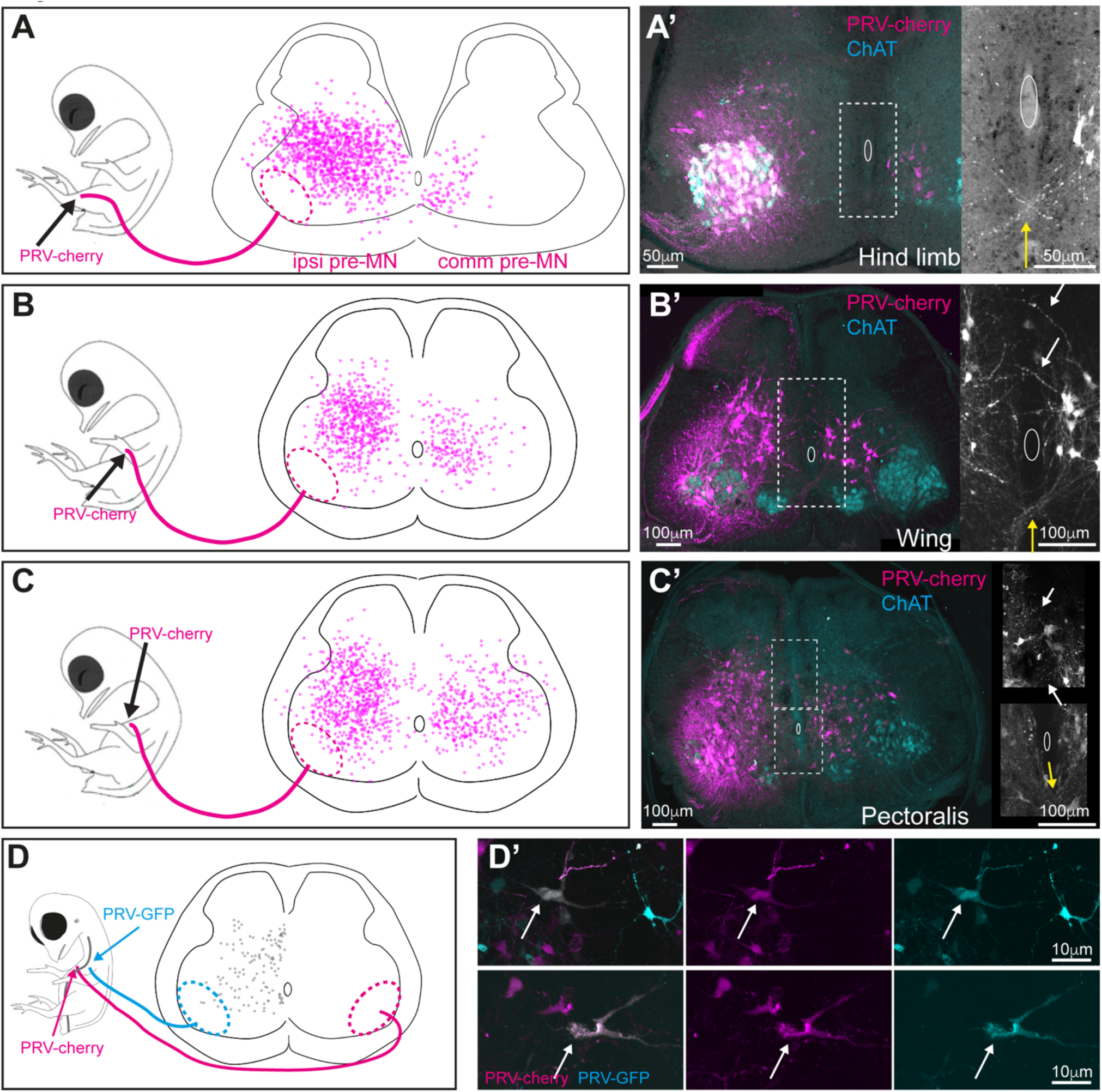
Patterns of pre-MNs in the brachial and lumbar chick spinal cord. **A-C.** Cell distribution maps in the brachial and lumbar segments of E15 spinal cord sections of chicks in which the hindlimb (A), distal wing (B), or pectoral (C) musculature were injected with PRV-cherry at E14. Data are from 1038 ipre-MNs and 88 cpre-MNs in A (3 embryos), 620 ipre-MNs and 210 cpre-MNs in B (4 embryos) and 734 ipre-MNs and 392 cpre-MNs in C (2 embryos). Ellipses in dashed magenta represent the areas occupied by motoneurons. A’-C’. Images of PRV-positive neurons supplemented with immunostaining for motoneurons (ChAT); right insets are enlargements of the midline boxed area showing axonal decussation of pre-MNs at the FP (yellow arrows) and RP (white arrows); white ellipses represent the central canal. **D.** Identification of a subpopulation of pre-MNs that innervate pectoral muscle motoneurons bilaterally. PRV-GFP and PRV-cherry were injected bilaterally at E14 into the pectoral muscles. Cell distribution map of bilaterally innervating pectoral pre-MNs seen at E15 is shown in gray. D’. Examples of bilaterally innervating pre-MNs; left panels: merged images; middle panels: PRV- cherry; right panels: PRV-GFP. Out of 500 and 730 cpre-MNs on each side (18 sections), 11.9% ± 0.05 and 10.9% ± 0.04 were double labeled.

At the lumbar sciatic level, axonal decussation was apparent only through the FP (Fig. 5D- E), with no axons crossing the GB. At the lumbar crural level, which is devoid of GB, dorsal midline crossing was apparent, with 31±12% of crossing fibers occurring through the RP (Fig. 5F,G, crural). Notably, the sciatic nerve innervates muscles of the shank and foot, while nerves originating from the crural plexus innervate the thigh muscles (*41, 42*). Importantly, in the swing phase of birds, the ankle flexion leads to the elevation of the feet, while the knee is relatively stable. Hence, the sciatic nerve mediates the flexion and extension of the ankle and foot joints during stepping. Significant dorsal midline decussation was also apparent at the cervical, brachial and thoracic levels (Fig. 5G), as well as sacral levels (not shown), suggesting that the GB serves as a physical barrier for axonal decussation at the sciatic plexus level, thus allowing hindlimb alternation. In contrast, dorsal midline crossing at brachial levels correlates with synchronous wing (forelimb) movements, similar to that seen in *hop* mice.

### Connectivity of pre-MNs in the chick spinal cord

The synchronous gait of *hop* mice results from ectopic midline crossing of excitatory pre-MN through the FP and RP (Fig. 1B) (*9, 43*). To analyze the patterns of connectivity of pre-MNs at the lumbar and brachial levels of the chick, we injected the trans-synaptic PRV virus (*44, 45*) into the leg, the distal wing musculature, and the main flight muscle, the pectoralis major. We observed substantial differences in the densities and laminar distributions of commissural pre-MNs (cpre- MNs) at lumbar versus brachial levels (Fig. 6A-C, right side). cpre-MNs accounted for 7.81% of labeled cells at the sciatic level, and for 25.3% and 34.8% of brachial level cells labeled from the distal wing and the pectoral muscle, respectively.

The distribution of ipsilateral pre-MNs (ipre-MNs) was similar at lumbar and brachial levels, occupying the medial spinal cord (Fig. 6A-C, left side; Fig. S6). In contrast, the distribution of cpre- MNs was markedly different across levels. At lumbar levels (both sciatic and crural) we observed cpre-MNs ventro-medially, with most cells ventral to the central canal (Fig. 6A, right side; Fig. S6). At the brachial level, cpre-MNs of the distal wing musculature and the pectoral muscle were distributed ventrally and dorsally to the central canal (Fig. 6B-C, right side). Likewise, decussation of pre-MNs differed markedly between lumbar and brachial levels, with axons crossing only the FP at the lumbar level (Fig. 6A’), but both the RP and the FP at the brachial level (Fig. 6B’-C’).

Synchronous gait in mice *hop* mutants is also associated with pre-MNs that innervate motoneurons bilaterally (*43*). To directly test whether the chick brachial cord contains such cells, we injected PRV-GFP and PRV-cherry into the left and right pectoral muscles, respectively (Fig. 6D, left). 10.9% of pectoral cpre-MNs were double-labelled, most located above the central canal and adjacent to the dorsal midline (Fig. 6D), with neurite processes oriented toward the dorsal midline (Fig. 6D’). Hence, like in *hop* mice, a subpopulation of the brachial pre-MNs innervate motoneurons bilaterally.

To examine the neurotransmitter phenotype of spinal pre-MNs, we processed sections from hindlimb and wing PRV-injected embryos for *in situ* hybridization with a marker for excitatory neurons (vGlut2). At the lumbar level, 46.5% of the ipre-MNs and 38.6% of the cpre-MNs were excitatory. At the brachial level, 35% of the ipre-MNs and 38% of the ventral cpre-MNs were excitatory. Of the brachial-specific cpre-MNs that reside dorsal the central canal, 47% were excitatory (Table S1). The chick brachial-specific pre-MNs are thus comparable to the ectopic cpre- MNs in *hop* mutants in location, wiring and excitatory phenotype (*14, 43*).

### Exogenous ephrin-B3 expression in the chick RP prevents midline axonal crossing

Our observations so far suggest that dorsal midline crossing and the wiring pattern of brachial pre- MNs in chick spinal cord might be linked to a lack of ephrin-B3 expression due to the genomic loss. To test this hypothesis, we examined whether expression of mouse ephrin-B3 in the chick would affect RP neurite crossing. Mouse ephrin-B3 or GFP, cloned into a Cre-dependent expression vector under the RP-specific Wnt1 enhancer element (*46*), were co-electroporated into the E3 chick spinal cord (Fig. 7A), and the extent of RP neurite crossing analyzed at E15. To exclude possible effects of exogenous ephrin-B3 in reverse signaling, we also used an ephrin-B3 isoform that lacks the intracellular domain (ephrin-B3-ΔC). Ephrin-B3 expression was confined to the RP (Fig. 7B), and the number of midline crossing neurites was significantly lower for ephrin-B3 (7.1 ± 3.1) or ephrin-B3-ΔC (7.1 ± 2.6) than for GFP (16.5 ± 4.6) (Fig. 7C-E). Hence, as in mammals, midline ephrin-B3 expression is sufficient to prevent midline crossing.

**Figure 7:**
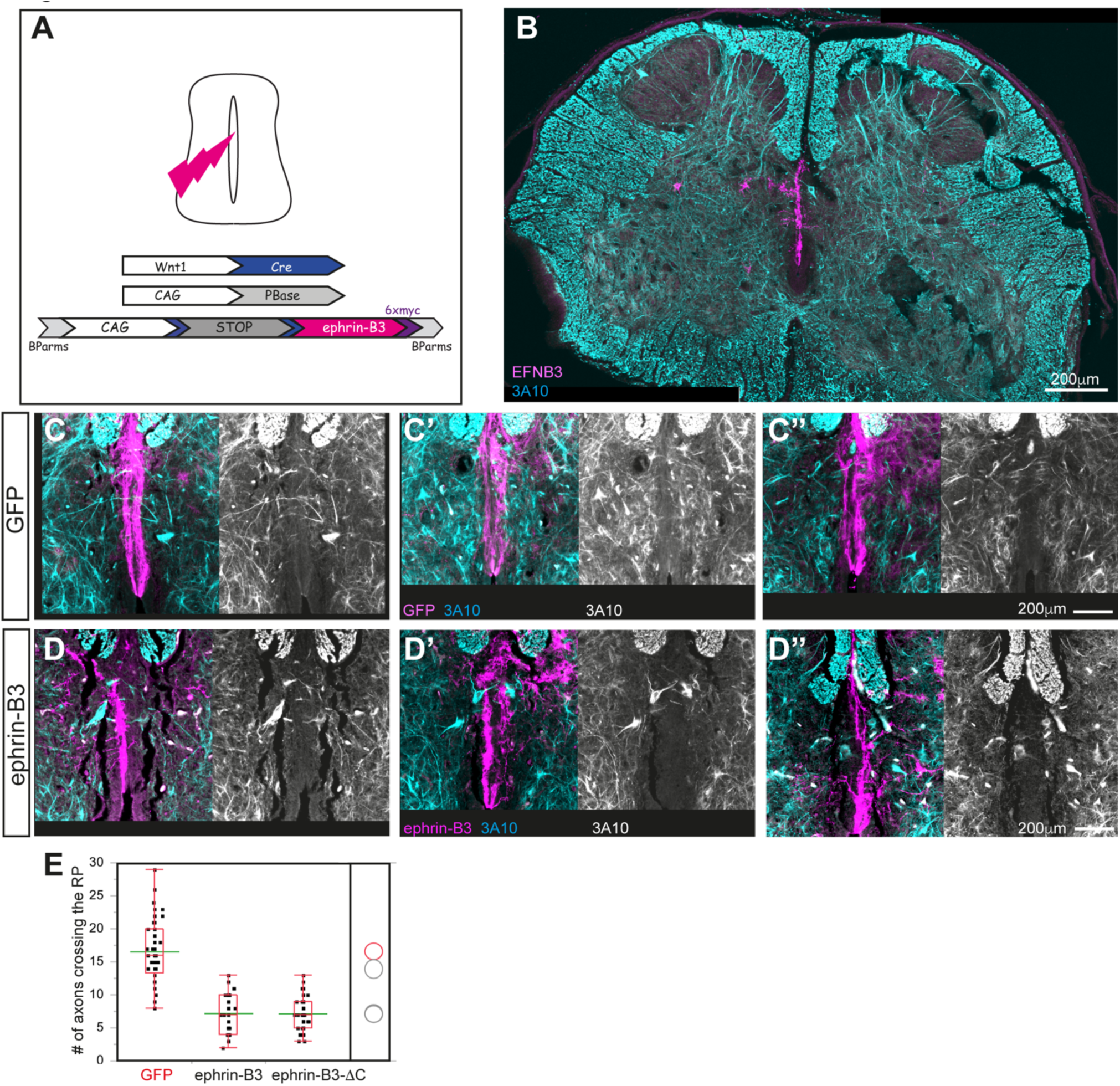
Expression of exogenous ephrin-B3 in the chick RP prevents midline crossing. **A.** Strategy for ectopic expression of the mouse ephrin-B3 at the roof plate. **B.** Cross-section from brachial spinal cord of E17 embryo expressing ephrin-B3 (magenta) at the dorsal midline, immunostained for neurofilament (cyan). **C,D.** Magnifications of 3 examples of dorsal midline region of spinal cords electroporated with GFP (C,C’,C”) or ephrin-B3 (D,D’,D”) (both in magenta), immunostained for neurofilament (cyan). The black and white images are the neurofilament (3A10) staining. Some neurites cross the GFP-expressing RP (C) but not the ephrin-B3-expressing RP. **E.** Quantification of dorsal midline crossing in spinal cords expressing GFP (N=43 sections, 3 embryos), mouse ephrin-B3 (N=71 sections, 3 embryos) or ephrin-B3-ΔC (N=59 sections, 3 embryos). The numbers of neurites crossing ephrin-B3- or ephrin-B3-ΔC-expressing RPs were significantly lower than in GFP-expressing controls (See supplementary statistics).

Next, after targeting ephrin-B3 expression to the RP at E3, PRV was injected into the wing musculature at E14 (Fig. 8A), and the numbers and distribution of cpre-MNs examined at E15.5. In controls, both ipre-MNs and cpre-MNs were seen, the latter both dorsal and ventral to the central canal (Fig. 8B), and their neurites crossed at both the FP and RP (Fig. 8B’). In ephrin-B3-expressing embryos, fewer cpre-MNs were seen, most ventral to the central canal (Fig. 8C,C’,C’’), and most neurites were observed to cross the FP (Fig. 8C,C’,C’’, right insets). Short cell extensions, possibly dendrites originating from ipre-MNs, did cross the RP (Fig. 8C, C’, right insets). We note that *in ovo* electroporation results in mosaics where only a portion of the cells express the transgene. Our efficiencies varied from 5% to 64% (sections with midline expression of ephrin-B3), thus some axons may circumvent ephrin-B3 and cross the midline close to non-EFNB3 expressing RP cells (Supp movie).

**Figure 8:**
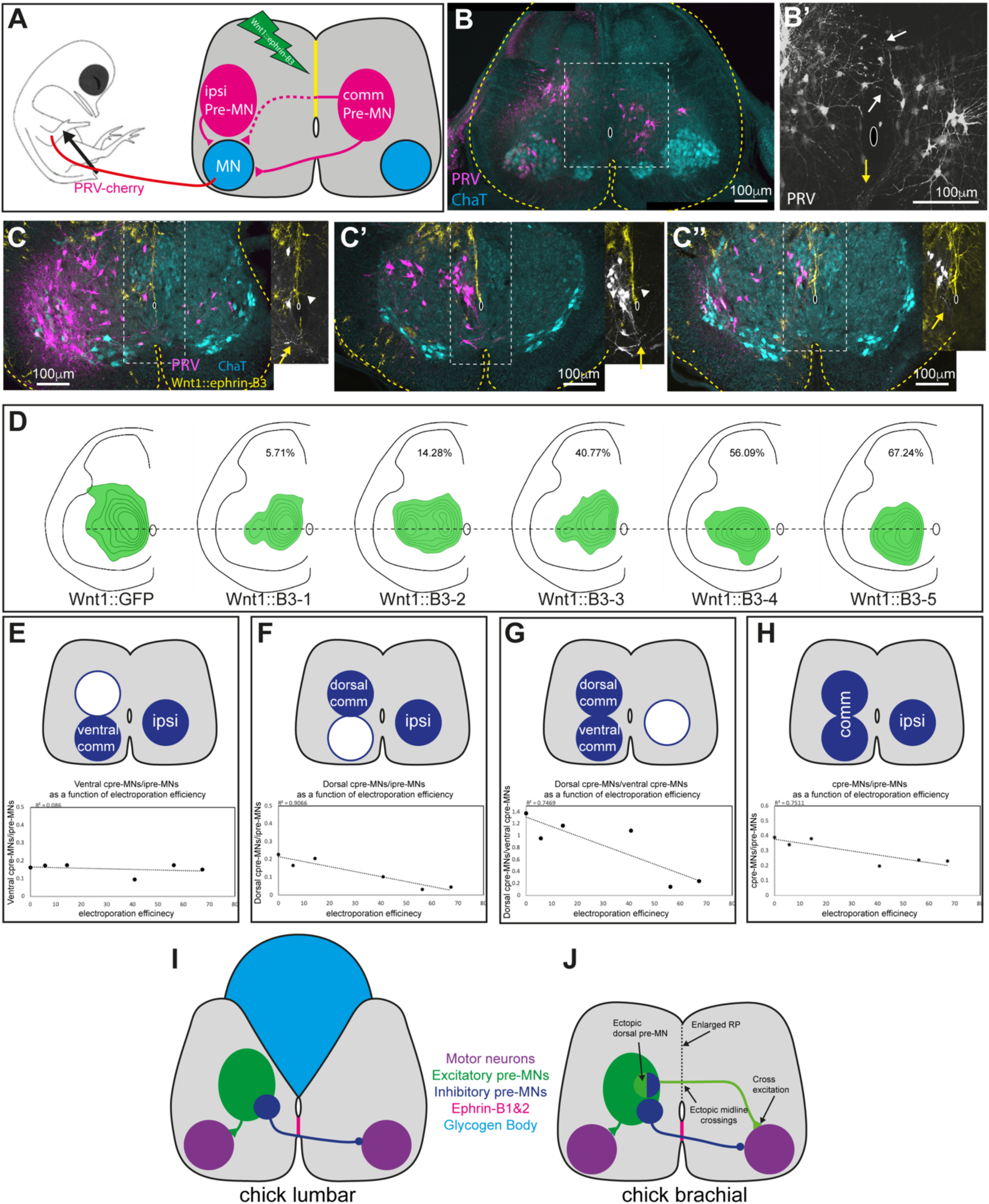
Expression of exogenous ephrin-B3 in the chick RP prevents pre-MN crossing. **A.** Schematic of the experimental setup. GFP or ephrin-B3 were expressed in the midline at E3. At E14, PRV-cherry was unilaterally injected into the wing musculature, and embryos fixed and processed 38 hr later. ChAT-immunostained (cyan) motoneurons (MN)and PRV-labeled (magenta) pre-MNs are indicated; dashed line represents cpre-MN axons that cross the RP. **B,C.** Transverse sections of control (B) and Wnt1::ephrin-B3-expressing E15 embryos (3 examples in C,C’C”). The black and white images are in B’ and the insets in C,C’,C” are PRV staining. In the control (B), cpre- MNs somata are located ventral and dorsal to the central canal and neurites of cpre-MNs are apparent at the FP (yellow arrows in B’) and RP (white arrows in B’). In Wnt1::mEFNB3 embryos (C, C’, C’’), the somata of pre-MNs are mainly located ventral to the central canal. Axons crossing the FP (yellow arrows) and few short neurites (white arrowheads) crossing the dorsal midline are shown in the right insets. **D.** Density maps of brachial cpre-MNs in control (GFP, N = 880 cells from six embryos) and five ephrin-B3-expressing embryos (N = 242, 467, 366, 49 and 67 cells). The fraction of sections containing ephrin-B3 expression along the entire dorsal midline is indicated at the upper part of each spinal cord. A ventral shift of the distribution is apparent in the high efficiency electroporated spinal cords of wnt1-B3-4 and wnt-1-B3-5. **E-H.** Plots demonstrating the correlation between ventral commissural/ipsilateral, commissural/Ipsilateral, dorsal commissural/ipsilateral and dorsal commissural/ ventral commissural pre-MNs ratios to the electroporation efficiency (as described in D), respectively. **I,J.** Schematic illustration of the circuitry at the lumbar (I) and brachial (J) avian spinal cord. We propose that dorsal midline crossing is prevented at the lumbar level by the glycogen body, while dorsal midline crossing at the brachial level is enabled due to the loss of ephrin-B3 activity. The brachial avian spinal cord shares significant morphological and wiring similarities with mouse *hop* mutants.

Next, we analyzed the spatial distribution of cpre-MNs in five embryos with variable efficiencies of wnt1::ephrin-B3 expression (Fig. 8D). A dorsal to ventral shift in the location of cpre- MNs is apparent when comparing the embryos with lower vs higher exogenous ephrin-B3 expression in the RP (1-3 vs 4-5 in Fig. 8D), suggesting a reduction in the size of the brachial- specific dorsal population of cpre-MNs. To further quantify this observation, we subdivided cpre- MNs into ventral and dorsal groups, based on location relative to the central canal, and examined their numbers compared to ipre-MNs. The ratios between pre-MNs populating the different zones were calculated and correlated with the efficiencies of ectopic expression of ephrin-B3, using linear regression (Fig. 8E-H). The ratio of ventral cpre-MNs to ipre-MNs was similar in controls and all experimental embryos (Fig. 8E), thus ephrin-B3 RP expression did not affect this population. The ratio of dorsal cpre-MNs to ipre-MNs was reduced in proportion to higher ephrin-B3 expression (Fig. 8F), as was their ratio to ventral cpre-MNs (Fig. 8G). The ratio of all cpre-MNs to the ipre-MNs was also negatively correlated with increasing ephrin-B3 expression (Fig. 8H). We conclude that robust ephrin-B3 RP expression results in fewer cpre-MNs that cross through the RP.

## Discussion

During vertebrate evolution, most terrestrial species adopted alternating limb motion as an effective gait strategy, while adaptation to powered flight in birds emerged via morphological changes that patterned wings from forelimbs, and required transition from alternate gait to synchronous flapping. Our genomic analysis indicates that the gene encoding the midline axonal repellent ephrin-B3 is missing in chicken and other Galliformes. In songbirds, the EFNB3 gene encodes an ephrin-B3 protein that does not bind to the EphA4 receptor, and whose expression in the spinal cord is reduced along the roof plate, probably due to the loss of a downstream enhancer. Morphological and circuit analyses of the chick spinal cord revealed similarities at the brachial level with mutant mice for components of the ephrin-B3/EphA4 pathway, while at the lumbar level it resembled wt mice (Fig. 8I,J). Importantly, dorsal midline expression of exogenous (rodent) ephrin-B3 in the chick brachial spinal cord prevents crossing of cpre-MN axons. Altogether, our findings support the notion that a reduction in ephrin-B3 expression and/or function plays an important role in shaping the wing level circuitry in the avian spinal cord.

### Loss of ephrin-B3 function in birds

Changes in gene copy number, enhancers, and protein sequence, arguably some of the foremost molecular events in evolution, are thought to have influenced locomotion patterns in vertebrates. For example, a point mutation that leads to a premature stop codon in *DMRT3* enabled Icelandic horses to develop their characteristic Tölt ambling gait (*47*). A modification and subsequent loss of a regulatory element of *GDF6* during the transition to bipedalism in humans is thought to have played a role in foot digit shortening (*48*). Limb loss in snakes has been shown to be a result of mutations in a limb-specific enhancer of Sonic hedgehog (*49*). Loss of function for ephrin-B3 leads to midline crossing in the spinal cord of rodents, resulting in a remarkable hopping gait (*9, 10*). Our findings now indicate that the ephrin-B3 gene has undergone significant modifications in birds compared to non-avian groups, likely impacting flight control circuits in the avian spinal cord.

Absence of *DNAH2* is seen in all birds where this genomic region could be assessed, suggesting that the loss of a putative ephrin-B3 enhancer was an early event in avian evolution, or possibly even in dinosaurs. The apparent loss of EFNB3 in chick and other Galliformes was likely a more recent event specific to that basal clade, since EFNB3 is present in Anseriformes and in several Neoaves, including the more recently evolved Passeriformes. It is hard to estimate the timing of mutations in the coding region, since high quality sequence is limited to a few species. Nonetheless, similar protein changes compared to mammals and non-avian saurospids are present in eagles and in songbirds (Fig. S2), which originated ∼65 and ∼45 million years ago, respectively (*16*). High-quality sequences of this region in the most basal avian clade, Palaeognathae, currently not available, will be critical for better understanding the timing and extent of avian EFNB3 changes.

In zebra finch, besides a predicted lack of reverse signaling due to loss of specific domains, we provide clear evidence that forward signaling through the EphA4 receptor is impaired. The lack of binding to avian EphA4, for example, is likely related to substitutions in residues 108-111 (in the mouse protein). This region is within the G 1-strand which forms a loop with the H 1-strand, and is part of the ligand-receptor dimerization interphase (*50*). The D108Q, L109R and L111V substitutions may therefore affect receptor binding by disrupting the formation of the G-H loop. Interestingly, ephrin-B3 is also a known receptor for the Nipah paramyxovirus (*Henipavirus* group) and the G-H loop is critical for infectivity (*51*), as shown by mutations in the H 1-strand of ephrin- B3 (*52*), suggesting possible related changes in spinal circuitry and viral susceptibility mechanisms.

### Phenotype of avian and *hop* mouse spinal cords

Our data show that the brachial spinal cord of the chick, which lacks an EFNB3 gene, shares similarities with mouse *hop* mutants in its morphology, and in the wiring and excitatory/inhibitory balance of cpre-MNs (Fig. 1B, 8J). The significance of the dorsal midline expansion, anatomically the most profound alteration in *hop* mutants, is unclear. While synchronous motoneuron activity is seen in the spinal cord of *hop* mutants using an *in vitro* fictive locomotion assay, severing of the dorsal midline does not restore left-right alternation (*8, 53*), arguing that bilateral coupling is facilitated by ventral midline crossing. However, *in vivo* disruption of the roof plate integrity is sufficient to induce synchronous gait in mice (*13, 14*). It is conceivable that in *hop* mutants, proprioceptive signals from weight-load receptors, which are used only *in vivo*, inhibit the bilateral coupling mediated by the dorsally crossing axons. In support of this hypothesis, removal of EphA4 from a subpopulation of dorsal interneurons results in ectopic dorsal midline crossing. Furthermore, the over-ground gait of these mice is normal, while during air-walking and swimming their limbs move in synchrony (*43*). In birds, dorsal midline crossing might be sufficient to instruct the bilateral coupling of the wings motoneurons, since weight load without ground support during flight does not affect the wings.

The importance of excitatory neurons for synchronous gait in mouse *hop* mutants is shown by conditional targeting of EphA4 in vGlut2 neurons, which results in hopping gait (*8*). Furthermore, a rather small shift in the excitatory/inhibitory balance in cpre-MNs, from 21% excitatory and 46% inhibitory in wt to 26% and 35% in the EphA4 null mouse, is sufficient to induce synchronous gait (*9, 53*). We found that the brachial dorsal population of cpre-MNs in chick is moderately but significantly more enriched (–47%) with excitatory neurons than in wt mice. Furthermore, the difference between this population and the brachial ventral and lumbar cpre- MNs, 38% and 35% of excitatory neurons, respectively (Fig. 6, Table S1), is similar to the difference between wt and *hop* mutant mouse. These observations support the notion that the brachial-specific population of cpre-MNs is sufficient to support synchronous activation of motoneurons involved in wing flapping.

Intriguingly*, t*he glycogen body (GB) (Fig. 8I) is a prominent but poorly understood dorsal lumbar gelatinous structure that arises from roof plate cells (*33, 34*) and is present across birds (*54, 55*). Our study has uncovered evidence that the GB serves as a major physical barrier to dorsal midline crossing at sciatic lumbar levels in the chick. We thus suggest that the GB is likely a major contributor to the alternating gait in birds, analogous to the molecular barriers to midline crossing in mice.

In summary, we provide evidence linking loss of ephrin-B3/ EphA4 signaling to the emergence of synchrony-mediating circuitry in the avian brachial spinal cord. While our findings do not exclude the importance of other molecular pathways, they suggest that modulation of ephrin-related mechanisms may have been an important contributor to the origin of flight in birds.

## Materials and Methods

### Genomics

The EFNB3 genomic region in birds has presented several difficulties for sequencing and assembling, likely due to very high GC content and density of poorly characterized repeat elements. The gene and its syntenic region are not present, or present only in partial form, in most Sanger- and Illumina-based avian assemblies, but are better represented in the more recent PacBio assemblies (Korlach et al., 2017). Furthermore, this region is likely part of a short avian micro-chromosome, which until the recent Swainson’s thrush genome had not been assembled and/or annotated in any species. Our current study of EFNB3 in birds has therefore required considerable search, annotation and curation efforts.

Briefly, we initially examined the annotated databases (NCBI’s RefSeq, Ensembl) available for >50 avian species, searching for gene predictions annotated as EFNB3 or ephrin B3-like, and using reciprocal BLAT alignments and verification of conserved synteny of the predictions to confirm their orthology with humans and non-avian vertebrates. We were able to identify EFNB3 and to verify its orthology in seven songbird species (including zebra finch), an eagle, and a parrot (kakapo). Notably, in the Swainson’s thrush EFNB3 and syntenic genes have been assigned to chr37, the first case any genes have been assigned to that avian micro-chromosome. We also note that two apparently identical zebra finch EFNB3 models (discontinued Gene IDs: 115492780 and 115492629) were present in an unplaced scaffold (NW_022045341.1) of an early PacBio assembly (bTaeGut1_v1.p; (*56*), but have been recently discontinued as this entire region is not present in the latest zebra finch Pacbio assemblies in NCBI. We used the discontinued XM_030261704.1 transcript prediction in our study, noting that it is well supported by ESTs and transcriptome data (Fig. S1A,B). Ensembl has EFNB3 predictions for the zebra finch, kakapo, and sparrowhawk, the latter in a short segment and right upstream of KDM6B. We also BLAST searched avian and non-avian RefSeq databases using as queries the genomic regions just downstream of avian gene predictions annotated as WRAP53 (or WD repeat containing antisense to TP53, a.k.a. telomerase Cajal body protein 1), or just upstream of avian gene predictions for KDM6B (or lysine specific demethylase 6B). This effort identified EFNB3 in an owl and in another songbird species, even though no EFNB3 predictions were present. We also BLAST searched all existing avian genomes (May 2020) as well as the respective WGS databases, using as queries the existing avian EFNB3 predictions, and manually examined all hits with scores >50, excluding cases that did not show top reciprocal alignments (typically other EFNB family members). This effort identified EFNB3 in a cormorant, and in a few other passerines and Anseriformes with limited synteny information, as well as in several species with incomplete data for synteny verification (a duck, a goose, several other passerines, eagles and cormorants, the seriema, a ruff, a crane, and a stork). As an EFNB3 was not present in the current Anna’s hummingbird PacBio assembly (the only such case for a non-Galliformes avian PacBio assembly), we also BLAST searched the pre-assembled reads (p- reads) for this assembly, and obtained evidence for an ortholog with similar synteny as in other bird groups. We also conducted BLAST searches for EFNB3 in the current assemblies of chicken (Pacbio-based GRCg6a) and quail (hybrid Coturnix japonica 2.1), as well as in the corresponding Pacbio p-reads and previous assemblies. These searches were extensive and permissive (>30 hit scores), followed by manual verification of detected hits. We also searched the BBSRC ChickEST Database and several brain transcriptome databases at E10, E17, P0 and 10 weeks old chick available in NCBI (ERR2576391, ERR2576392, ERR2576465, ERR2576464, ERR2576488, ERR2576584, ERR2576585, ERS2480079), but found no evidence of EFNB3.

### Animals

Fertilized White Leghorn chicken eggs (Gil-Guy Farm, Israel) were incubated in standard conditions at 38 °C. All experiments involved with animals were conducted in accordance with the designated Experiments in Animals Ethic Committee policies and under its approval. Experiments were performed on adult male and female mice. EphA4^lox^ mice (*57*) were crossed to PGK-Cre mice (*58*) to generate full knock-out of the EphA4 gene. Animals were kept and used in accordance with regulations from the government of Upper Bavaria.

### DNA

For the binding assay in COS cells, the mouse ephrin-B3 was cloned into pSecTag-B plasmid (Termo Fisher Scientific, Waltham, USA). Six repeats of the myc epitope were cloned at the carboxyl end of ephrin-B3. For cell surface detection of the mouse ephrin-B3, the kappa chain signal sequence plus 3 repeats of the myc epitope were cloned upstream to amino acid 30 of the mouse gene. The sequence of the full-length zebra finch ephrin-B3 transcript was reconstructed from RNA-seq reads (Fig. S1B,C) using the Trinity platform (*59*). The sequence matched the exonic sequence of the EFNB3 transcript prediction (XM_030261704.1). For cloning the zebra finch ephrin-B3, an optimized sequence (available upon request) of residues 21-171 (Fig. S2), predicted to encode the extracellular domain, was synthesized (ITD, Coralville, USA) and cloned downstream to an Ig kappa chain signal sequence in the pSecTag-B plasmid. The transmembrane and intracellular portions of the zebra finch transcript were obtained by fully sequencing the EST DV945855 cDNA clone. The synthetic 5’ portion of the gene plus the 3’ DV945855-derived sequence were fused using a PCR-generated EcoRV site at the synthetic 5’ part and an internal SmaI site in the 3’ DV945855-derived sequence. For detection of the *in vitro* transcribed protein, 4 repeats of the myc epitope were inserted between the Ig kappa chain signal sequence and predicted residue 21 of zebra finch ephrin-B3. The EphA4-AP fused gene was generated by cloning amino acids 1-543 of the chick EphA4 (*60*) into the pSecTag-B plasmid. The SEAP gene was cloned downstream to EphA4.

### Cell culture and transfection

COS-7 cells (ATCC) were cultured in DMEM media (Sigma, Saint Louis, USA) supplemented with 10% fetal bovine serum 100U/ml penicillin, 100 mg/ml streptomycin and 2 mM glutamine (Biological industries, Beit HaEmek, Israel). COS-7 cells were transiently transfected with appropriate cDNA using PEI (Polyethylenimine, linear, MW 25000; Polysciences, Inc).

### AP-fusion protein binding experiments

To produce secreted EphA4-AP (alkaline phosphatase, placental) fusion protein, COS-7 cells were transiently transfected with appropriate cDNA. Post-transfection cells were supplemented with Opti-MEM (Gibco), and media collected after 48 hr. The media was concentrated through a 30K ultracell filter (Merck Millipore, Tullagreen, Ireland). The amount of the AP fusion protein was quantified by SDS-PAGE analysis and Coomassie staining. The amount of EphA4-AP was calibrated by comparison to known amounts of EphA4-Fc fusion protein (R&D, Indinapolis, USA). AP activity was measured using para-nitrophenylphosphate substrate (pNPP, Sigma).

Binding assays were carried in COS-7 cells, transiently transfected with the mouse or zebra finch ephrin-B3. Following over-night (O/N) incubation, cells were incubated with varying amount of EphA4-AP in binding buffer (Hanks buffer saline supplemented with 20 mM Hepes, PH 7.3, 0.05% BSA, 5 mM CaCl2, 1 mM MgCl2 and 10 µg/ml heparin sodium salt (Sigma) at room temperature (RT) for 70 min. For visualization of AP staining, cells were plated on glass coverslip coated with poly-lysine (0.005%, Sigma), and following AP-ligand binding as described above, cells were washed with-binding buffer, fixed in 4% paraformaldehyde (PFA) in PBS for 15 min at RT, and then washed with HBS solution (20 mM HEPES pH 7.3, 150 mM NaCl). Endogenous AP activity was inactivated by incubation at 65 °C for 100 min. For color reaction, cells were incubated with NBT/BCIP substrates (Roche, Basel, Switzerland) in AP buffer (100 mM Tris pH 9.5, 100 mM NaCl, 5 mM MgCl_2_), O/N. After washing in HBS and fixation for 15 min in 4% PFA, and an additional washing in HBS, coverslips were mounted under mounting medium (Thermo Scientific, USA).

For colorimetric assays, following AP-ligand binding and washing, cells were incubated O/N at 4°C in lysis buffer (10 mM Tris, ph 8.0, with 1% Triton X100). The supernatant was subjected to heat inactivation for 15 min at 65°C. Lysates were transferred to 96-well high protein absorbance plate (Thermo Scientific, Nunc International USA) and the level of AP activity was determined following addition of equal volume of 2 mg/ml pNPP substrate in reaction buffer (200 mM Tris, pH 9.5, 10 mM MgCl2, 200 mM NaCl). Plates were incubated at RT for 30 min to 1 hr and optical density was determined at 405 nM using an ELISA plate reader.

### Ephrin-B3 surface staining

COS-7 cells were transiently transfected with the mouse or zebra finch myc-EphrinB3 expression vectors. 24 hr post-transfection, cells were incubated with mouse anti-myc (9E10) antibody in phosphate buffered saline (PBS), washed with PBS, and subsequently incubated with Rhodamine Red X (RRX) labeled anti-mouse antibody and washed again. Cells were fixed in 4% paraformaldehyde/0.1 M phosphate buffer and mounted before imaging.

### In-ovo electroporation

A DNA solution of 5 mg/mL was injected into the lumen of the neural tube at HH stage 17– 18 (E2.75-E3). Electroporation was performed using 3 × 50 ms pulses at 25–30 V, applied across the embryo using a 0.5 mm tungsten wire and a BTX electroporator (ECM 830). Following electroporation, 150–300 μL of antibiotic solution, containing 100 unit/mL penicillin in Hank’s Balanced Salt Solution (Biological Industry, Beit-Haemek) was added on top of the embryos. Embryos were incubated for 3–19 days prior to further treatment or analysis. Roof plate-specific expression was achieved utilizing the Wnt1 or DNAH2 enhancers, coupled with a dorsal electrode positioning.

### Immunohistochemistry and *in-situ* hybridization

Embryos were fixed overnight at 4°C in 4% paraformaldehyde/0.1 M phosphate buffer, washed twice with phosphate buffered saline (PBS), incubated in 30% sucrose/PBS for 24 h, and embedded in OCT (Scigen, Grandad, USA). Cryostat sections (20 μm) were collected on Superfrost Plus slides and kept at −20°C. For thicker sections, spinal cords were isolated from the fixed embryos and subsequently embedded in warm 5% agar (in PBS), and 100 μm sections (E12–E17) were cut with a Vibratome. Sections were collected in wells (free-floating technique) and processed for immunolabeling. The following primary antibodies were used: rabbit polyclonal anti-GFP 1:1000 (Molecular Probes, Eugene, Oregon, USA), mouse anti-GFP 1:100, goat anti-GFP 1:300 (Abcam, Cambridge, UK), rabbit anti-RFP 1:1000 (Acris, <u>Hiddenhausen, Germany</u>), goat anti-ChAT 1:300 (Cemicon, Temecula, CA, USA), mouse anti-neurofilament 1:100 (3A10), mouse anti-myc 1:10 (9E10), rabbit anti-myc 1:100 (Abcam). The following secondary antibodies were used: Alexa Fluor 488/647-AffiniPure Donkey anti-mouse, -rabbit and -goat (Jackson), and Rhodamin Red-X Donkey anti-mouse and -rabbit (Jackson). Images were taken under a microscope (Eclipse Ni; Nikon) with a digital camera (Zyla sCMOS; Andor) or a confocal microscope (FV1000; Olympus). *In situ* hybridization was performed as described (Avraham et al., 2010). The following probes were employed: chick ephrin-B1, ephrin-B2, EphA4 and Vglut2, zebra finch ephrin-B3 and mouse ephrin-B3. Probes were PCR-amplified from brain cDNAs of E6 chick embryo, P4 zebra finch, and E15 mouse, respectively, using the following primers. PCR products were verified by sequencing prior to use.

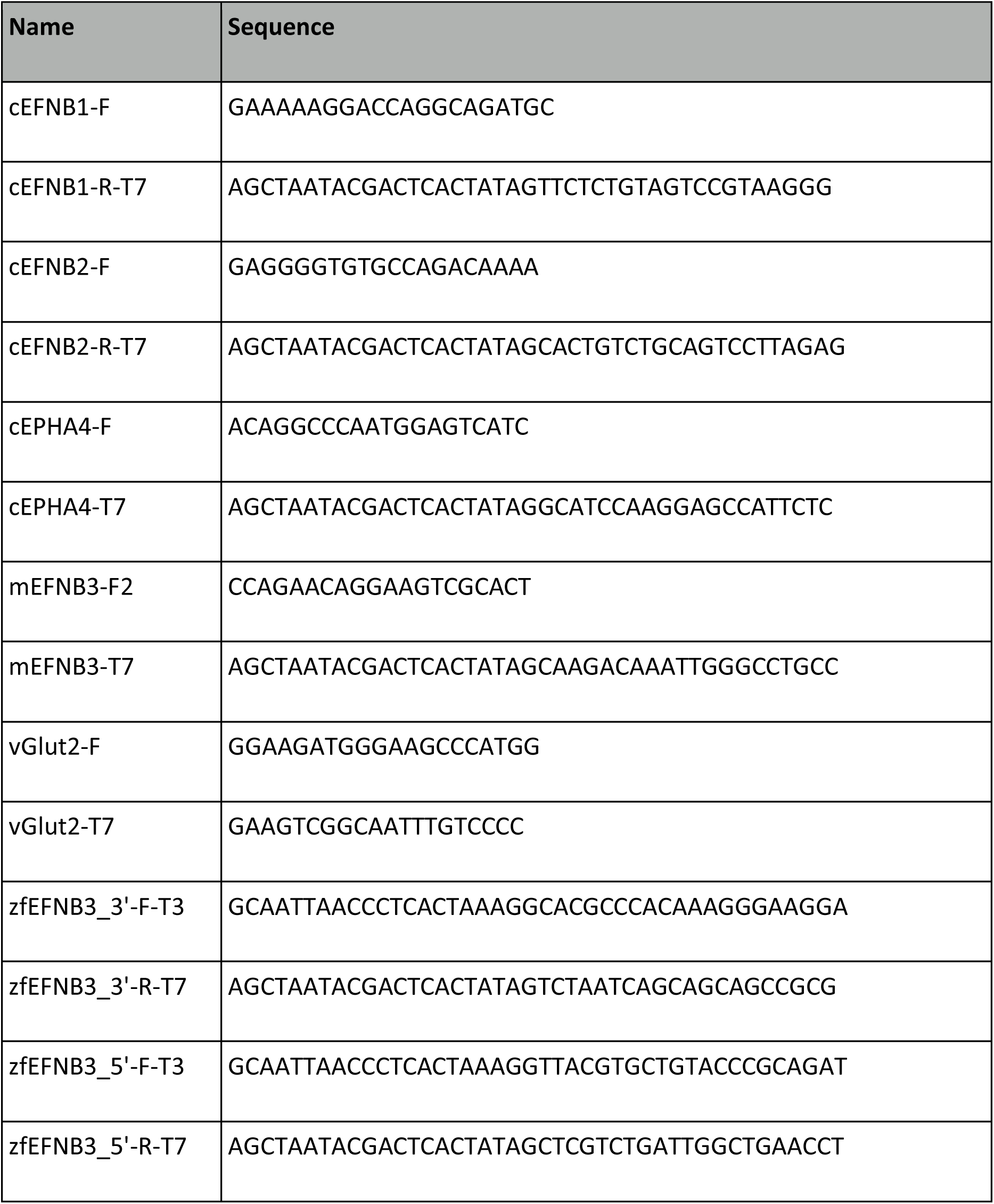

### PRV injection

We used two isogenic recombinants of an attenuated PRV strain (PRV Bartha) that express enhanced GFP (PRV152) and monomeric red fluorescent protein (PRV614). The viruses were harvested from Vero cell cultures at titers 4 × 108, 7 × 108 and 1 × 109 plaque forming units (pfu/mL), respectively. Viral stocks were stored at −80°C. For pre-motoneuron infection, injections of 3 μL of PRV152 or PRV614 were made into the thigh or wing musculature of E13 or E14 chick embryos, using Hamilton syringes (Hamilton; Reno, NV, USA) equipped with a 33-gauge needle. The embryos were incubated for 36–40 hr and sacrificed for analysis. For bilateral pre- motoneurons infection, PRV152 and PRV614 were each injected into the pectoralis musculature on opposite sides of the embryo midline.

### Cell distribution and density plots construction

The codes for the cell distribution and density plots imaging and analysis were written in Matlab. The distribution plots were constructed by calculating the relative midline expression, after background subtraction and a subsequent length normalization between all analyzed sections. The density plots were generated based on cross section images transformed to a standard form. The background was subtracted, and the cells were filtered automatically based on their soma area or using a manual approach. Subsequently, the neurons were depicted by semi-transparent circles or their distribution was visualized by two-dimensional kernel density estimation, using the MATLAB function “kde2d”. Unless indicated otherwise, a contour plot was drawn for density values between 20% and 80% of the estimated density range, in six contour lines. An additional script was used for pre-motoneuron classification based on their neurotransmitter identity and laminar location.

A similar density visualization procedure was done for the EphA4 density plots (Fig. 3E’ and F’), but was preceded by signal binarization and then an averaging filter (5 cycles) was used to achieve a smoother density signal. The intensity of EphA4 signal as a function of distance from the midline was obtained by summation of the binary signal over the horizontal axis, followed by a rescaling step for each image. Statistical analysis was done after subdividing the distance around the dorsal midline spanning from -15 to 15 (the relative distance between the central canal – 0, and the white/gray matter boundary – 100 and -100). into 6 equal intervals, and t-test was performed for the grouped data of each interval, originating from E10 and E16 data.

## Statistical Analysis

**Fig. 3I.**
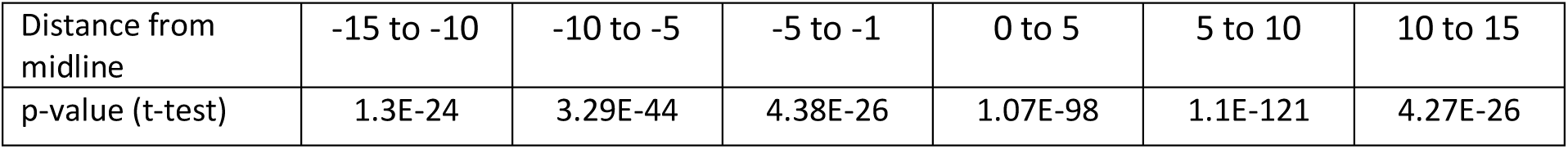

**Fig. 4G.**
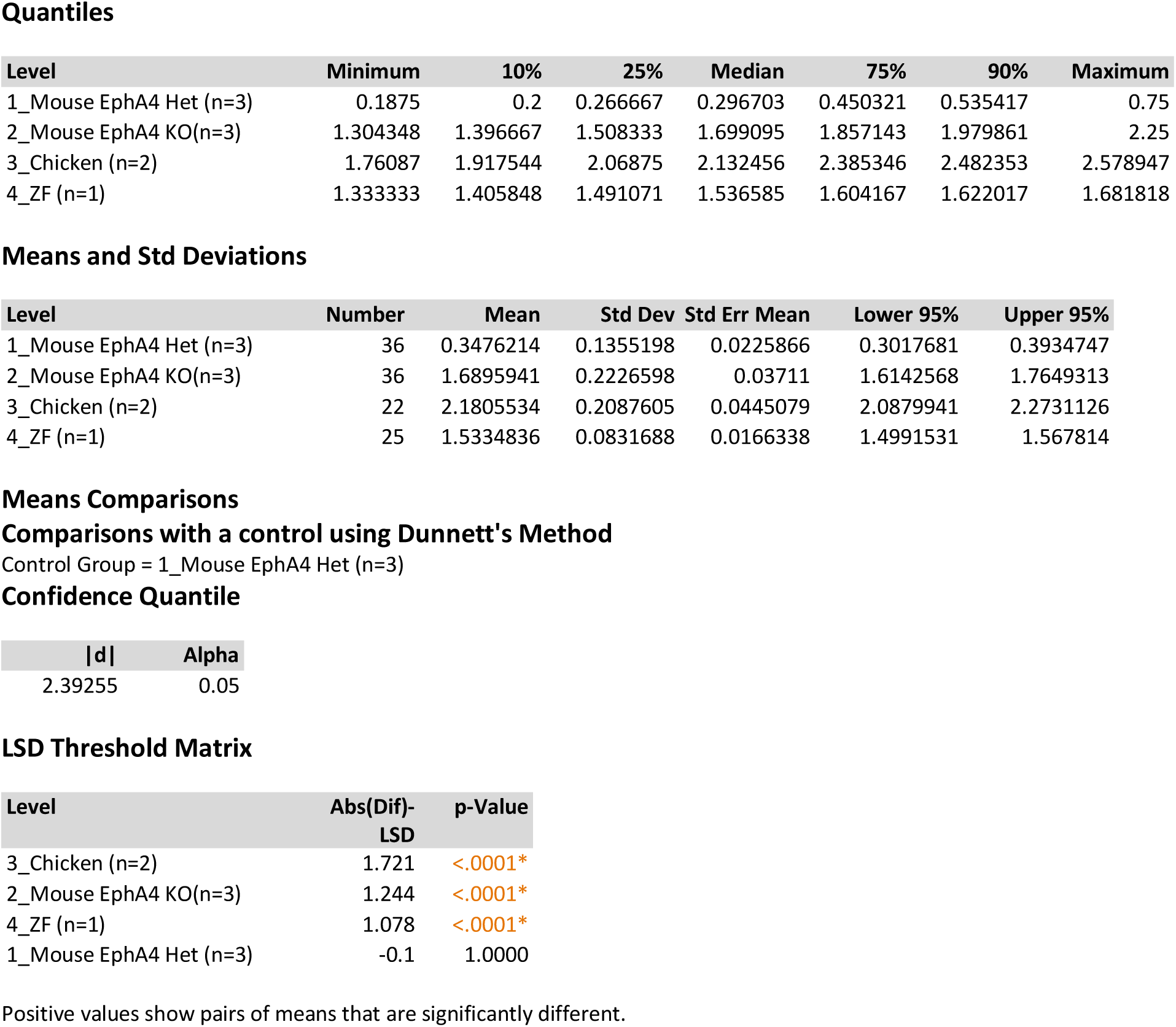

**Fig. 5G.**
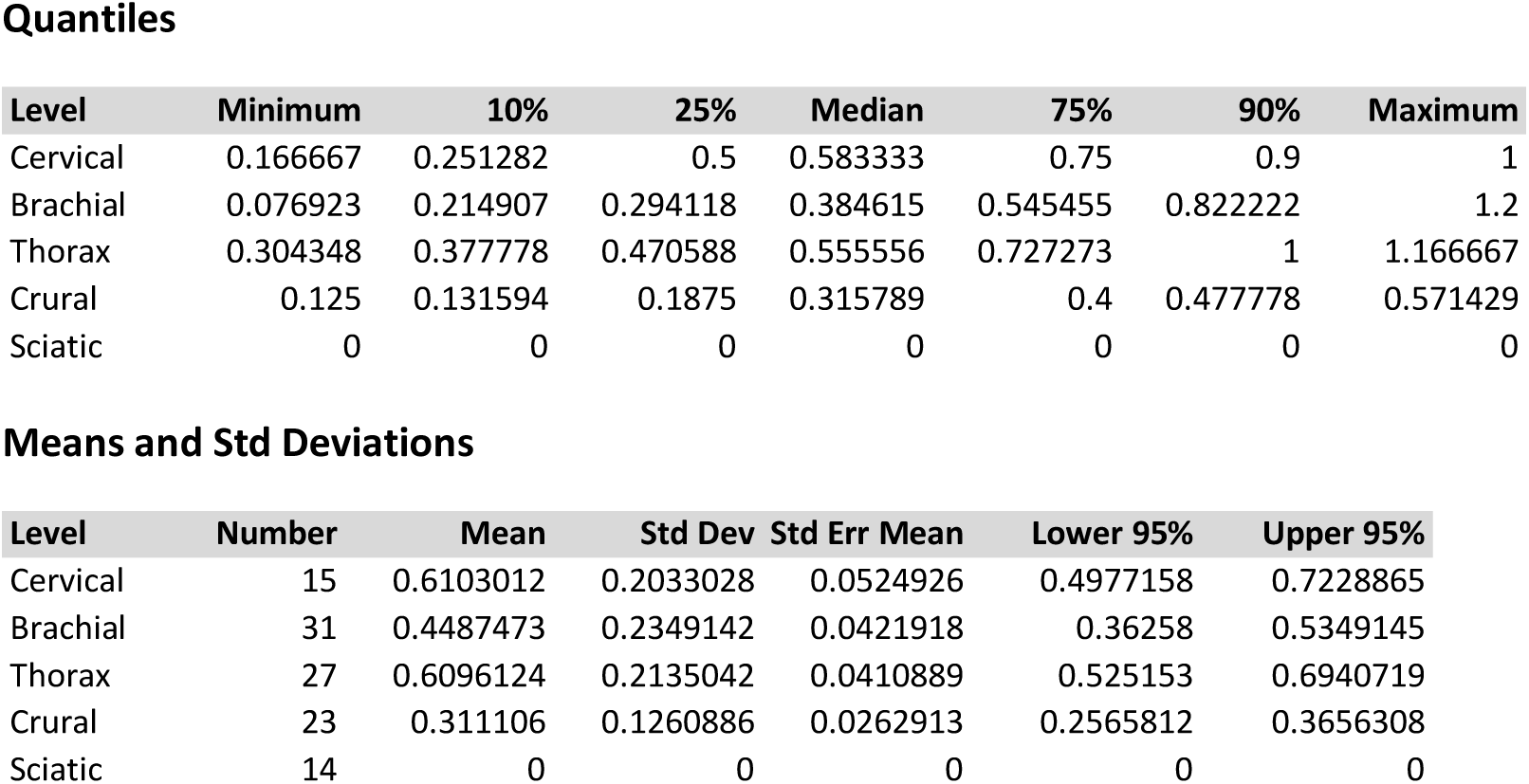

**Fig. 7E.**
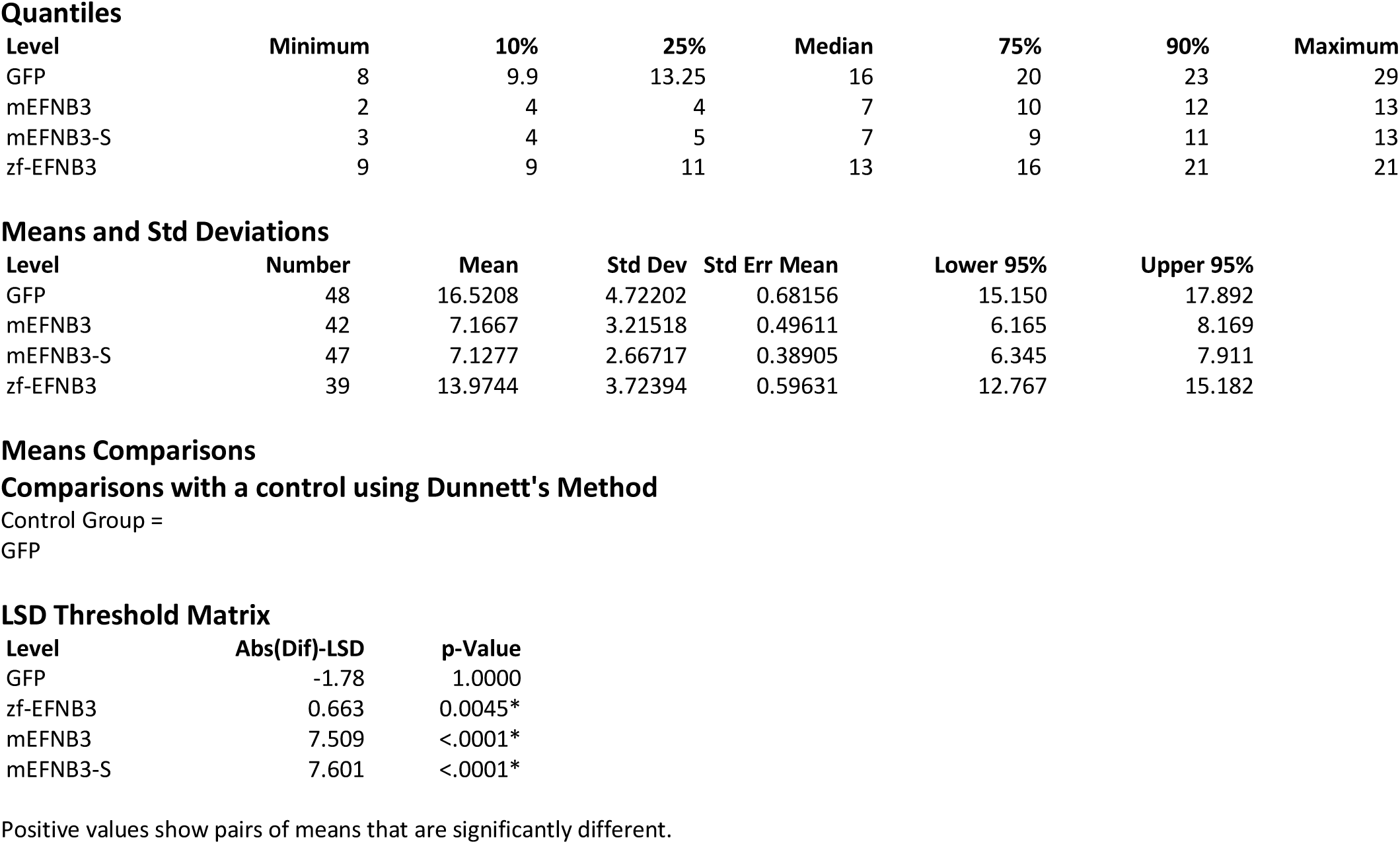

**8E-8H.**
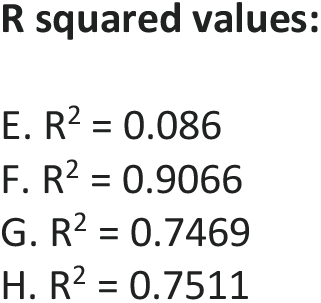

## Acknowledgments

The authors thank Tamir Lederman for technical assistance; Jerome Gros for providing the Pacbio p-reads of the quail genome, Wes Warren for providing the Pacbio p-reads of the chicken genome, and Anat Barnea for providing zebra finch hatchlings.

## Author contributions

AK, CVM, BH: research design, data analysis; BH: circuit analysis, in situ hybridization; OM: circuit analysis; RSM: enhancer analysis; JE: protein studies; SF, PVL: genomics and gene structure analysis; SP, RK: generated conditional ephrin-B3 knockout mouse; AK, CVM: wrote manuscript.

## Data and materials availability

The data was deposited at any archive.

## Supplementary Materials

**Fig. S1.**
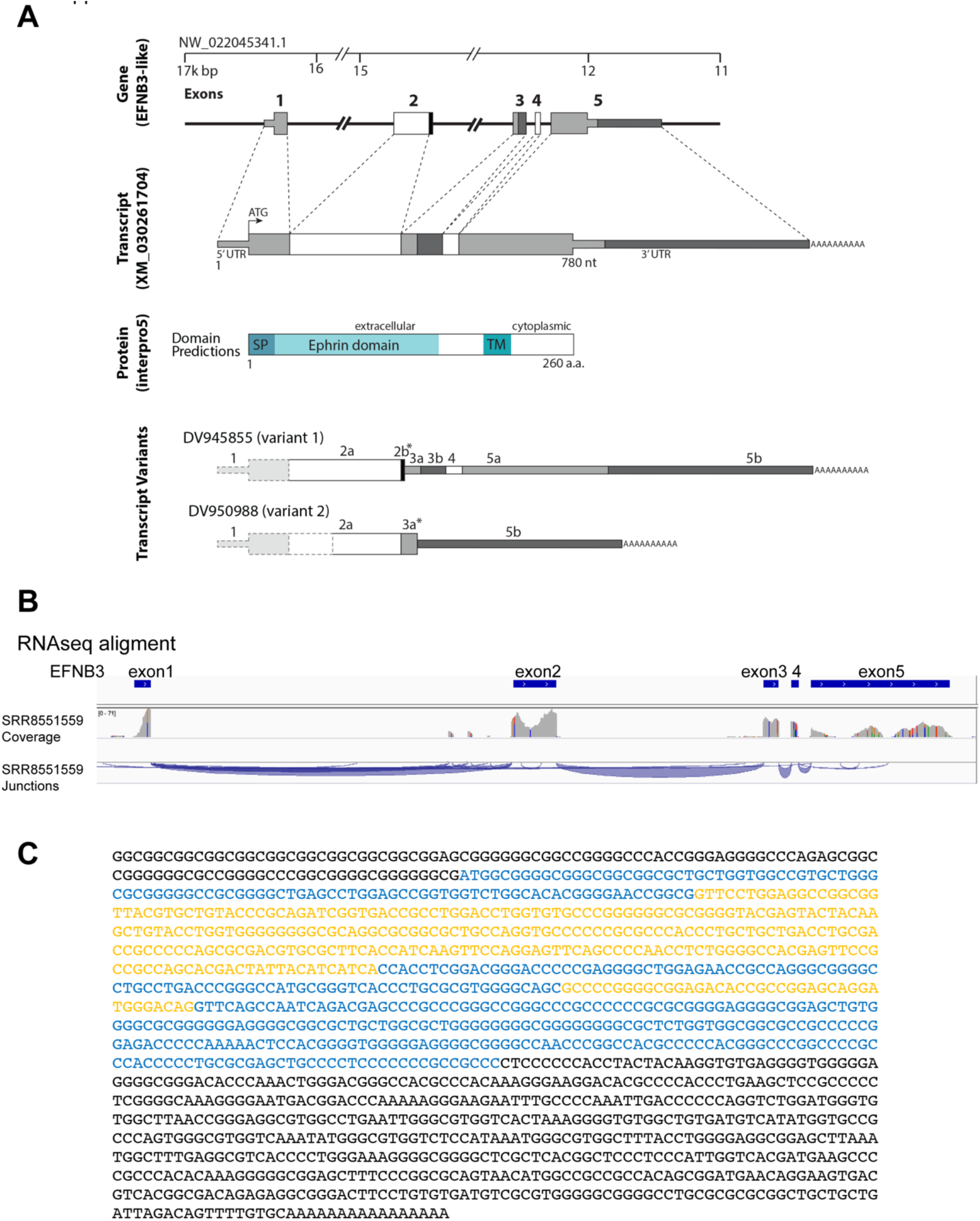
Ephrin-B3 gene and transcripts in zebra finch. A. Zebra finch ephrin-B3 gene, predicted transcript, predicted protein, and transcript variants. The two EST clones that include coding exons are DV945855 and DV950988. Both include frameshifts that result in premature *termination (indicated by asterisks)*. B. RNAseq reads of Ephrin-B3 in zebra finch. RNAseq data in zebra finches provide expression support for this structure and for expression of all five exons, and is consistent with the predicted polyA site based on multiple ESTs. C. The sequence of the full length zebra finch ephrin-B3 transcript was reconstructed from RNA- seq reads using the Trinity platform (*59*). The coding exons are indicated in alternating blue and yellow colors. The 5’UTR (black) is included in the 1^st^ exon and the 3’UTR (black) is included in the 5^th^ exon.

**Fig. S2.**
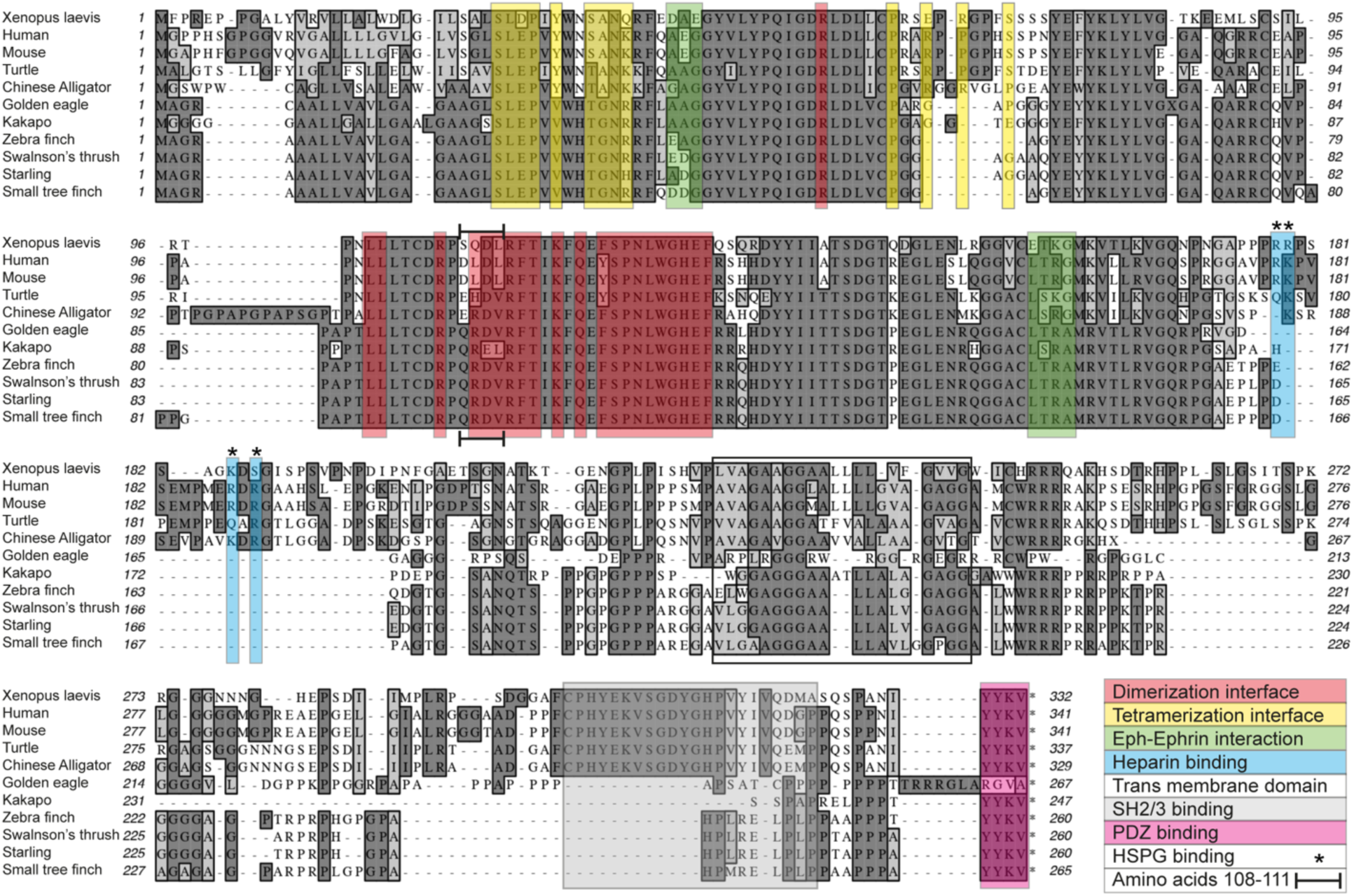
Phylogeny of the ephrin-B3 protein. Sequence alignment of the predicted ephrin-B3 proteins from different species. The T-Coffee algorithm was used for alignment of multiple sequences. Avian-specific deletions were detected in the heparin binding, tetramerization interface and SH2/3-binding domains (indicated by blue - with asterisks, yellow and gray rectangles, respectively), and specific mutations observed in the dimerization interface domain (indicated by red rectangles and horizontal bars in residues 108-111 of mouse). The open box indicates the transmembrane domain.

**Fig. S3.**
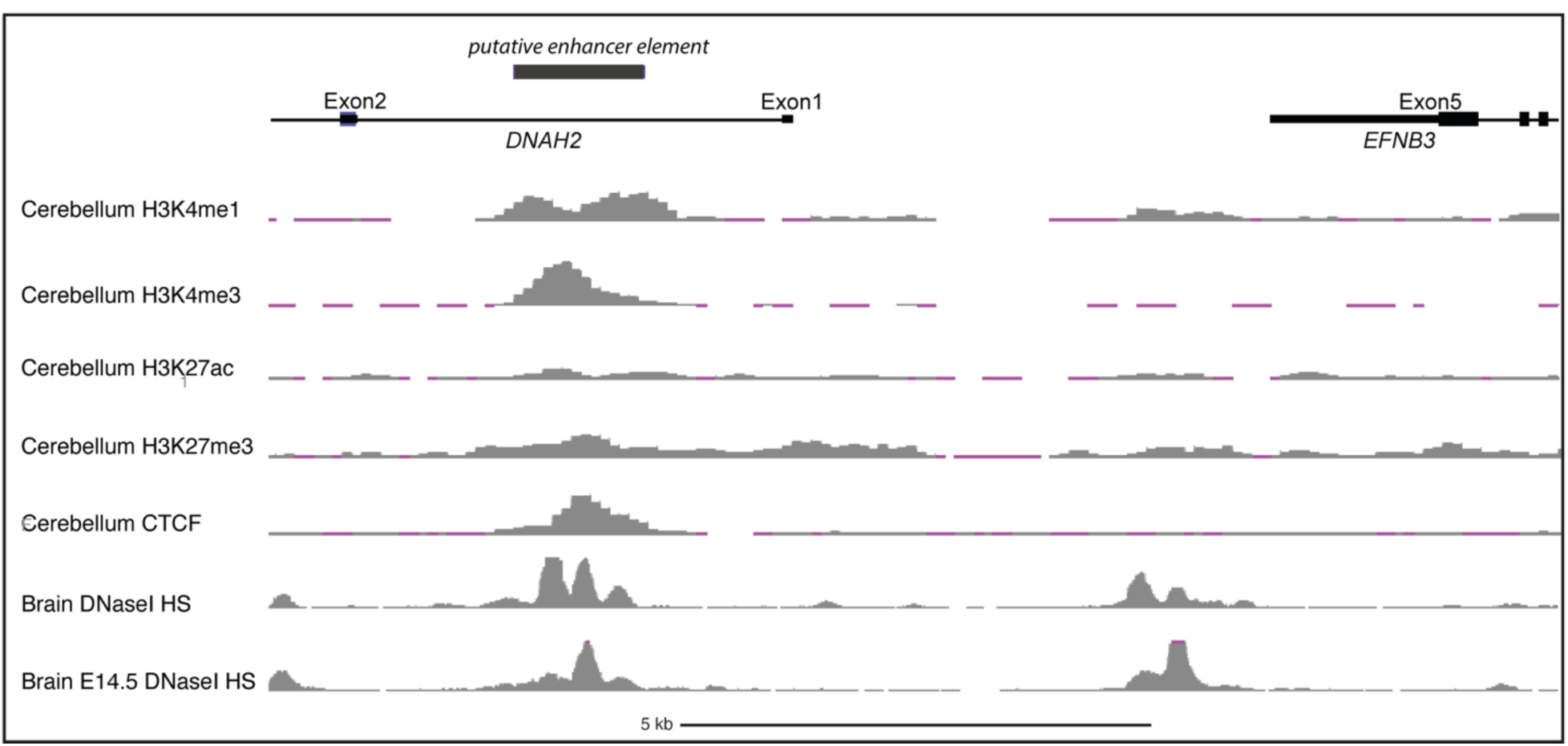
A putative EFNB3 enhancer in mouse DNAH2. A display of ENCODE analysis of candidate cis-Regulatory elements in mouse chr11:69,546,195- 69,547,544; shown is the region that includes EFNB3 (exons 3-5) and DNAH2 (exons 1-2) genes on the minus strand. A DNaseI hypersensitive (HS) site is in the 1^st^ intron of DNAH2 (chr11:69,546,195-69,547,544 in GRCm38/mm10). This motif binds modified histones (H3K4me3, H3K4me1 and H3K27ac) that are associated with active enhancers; histone H3K27me3 is associated with repression of developmental genes, as well as CTCF – a nuclear factor associated with active enhancers (*61, 62*). Binding of these proteins is apparent in the mouse embryonic cerebellum and brain. The data was obtained from UCSC genome browser (https://genome.ucsc.edu/index.html).

**Fig. S4.**
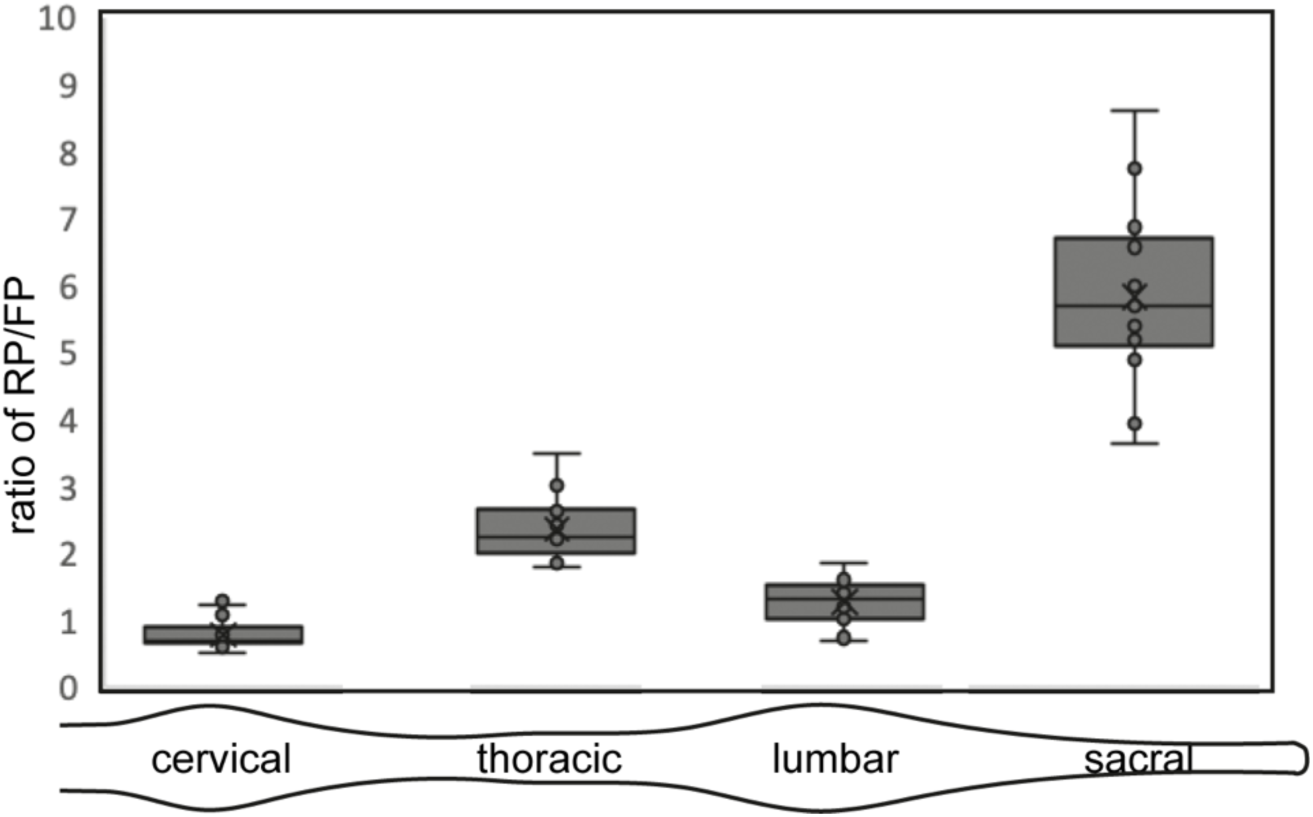
RP/FP ratio along the longitudinal axis of wt mice. The ratio between the roof plate and floor plate lengths at different segmental levels. The limb-innervating segments exhibit a lower RP/FP ratio compared to the thoracic and sacral levels. Analysis was done on images from the Mouse spinal cord atlas (Allen Institute for Brain Science; N = 17 mice, 15-16 section per mouse). Box plot representation of the roof plate (RP) to floor plate (FP) length ratios at the cervical, thoracic, lumbar and sacral levels at P4.

**Fig. S5.**
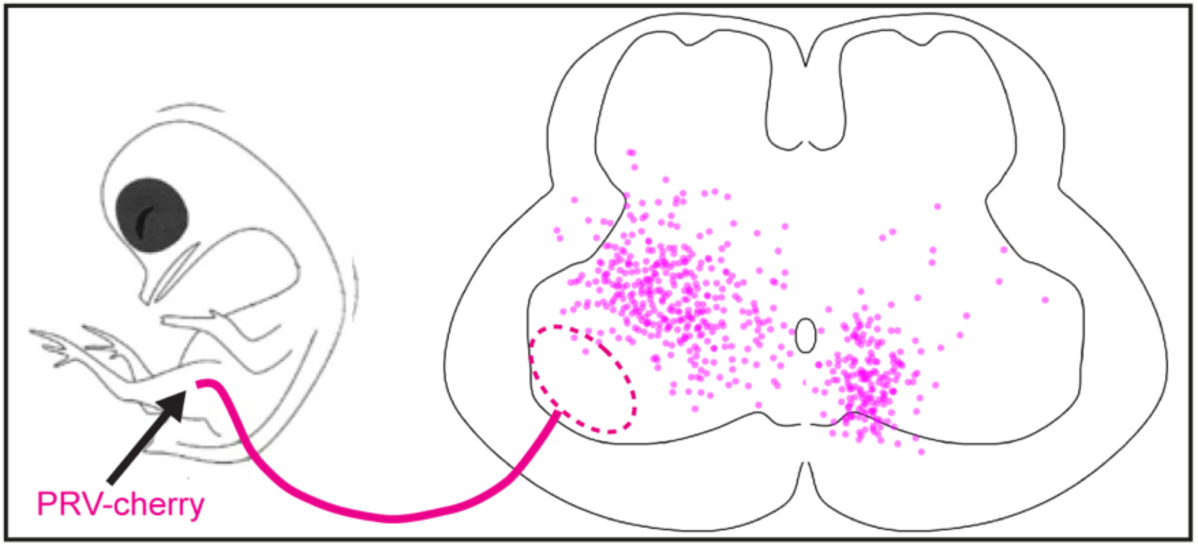
Distribution of pre-MNs at the spinal crural plexus level. pre-MN map at the crural lumbar level; shown are 361 ipre-MNs and 192 cpre-MNs, respectively (N=3 embryos).

**Fig. S6.**
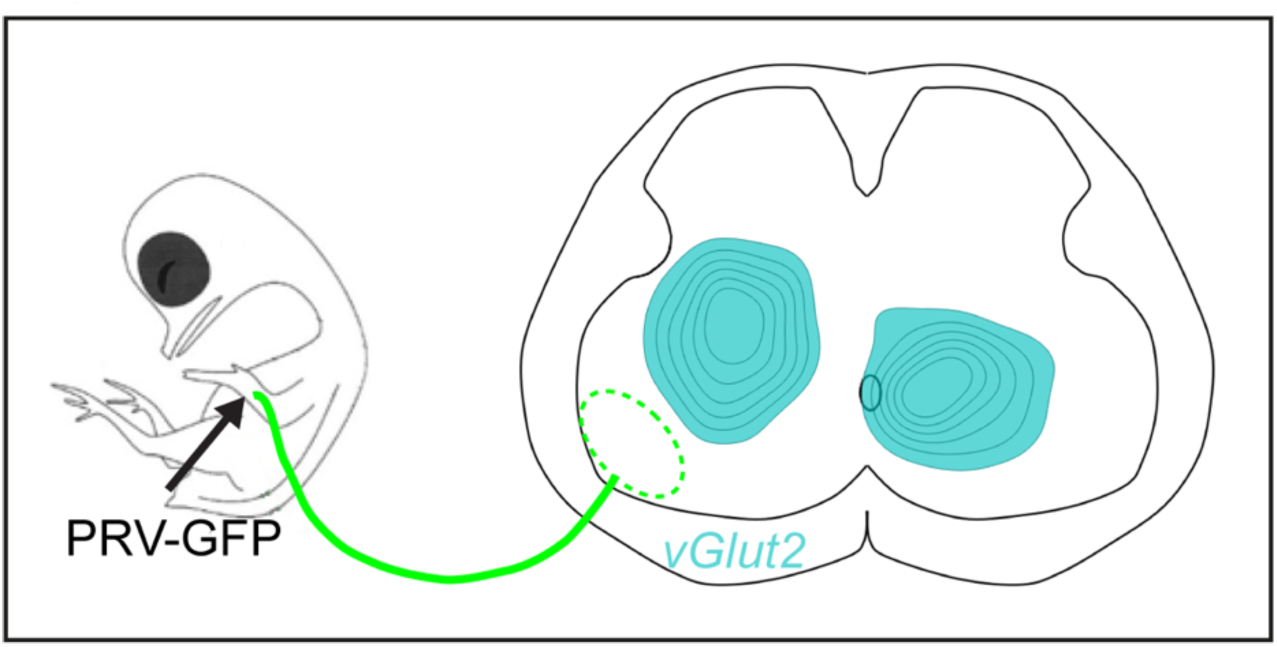
Brachial pre-MN neurotransmitter identity. Experimental design for brachial pre-MN labeling and *in situ* hybridization for VGlut2. Density maps of ipre-MNs and cpre-MNs are shown.

**Table S1.** The ratio of excitatory pre-MNs in the lumbar and brachial levels of the chick spinal cord.

| level | side | % vGlut2 | # |
| --- | --- | --- | --- |
| Lumbar sciatic | ipsi | 46.46 | 145 |
|  | comm | 38.62 | 145 |
| brachial | ipsi | 35 | 274 |
|  | Ventral comm | 38 | 29 |
|  | Dorsal comm | 47 | 51 |

